# scCoBench: Benchmarking single cell RNA-seq co-expression using promoter-reporter lines

**DOI:** 10.1101/2025.05.26.656221

**Authors:** Tran N. Chau, Kook Hui Ryu, Razan Alajoleen, Bastiaan O. R. Bargmann, John Schiefelbein, Song Li

## Abstract

Single-cell RNA sequencing (scRNA-seq) has become a powerful tool for uncovering transcriptomic heterogeneity and reconstructing gene regulatory networks in complex tissues. However, the sparsity, high noise levels, and dropout events inherent to scRNA-seq data pose challenges for accurate inference of gene-gene relationships. In this study scCoBench, we systematically benchmark correlation metrics, pseudo bulk analysis, and imputation methods using promoter-reporter and native gene pairs as internal controls to evaluate the performance of ten widely used gene-gene co-expression measurements. Interestingly, we found that commonly used data scaling and normalization approaches lead to lower correlation between promoter reporter and native gene pairs in most of the co-expression methods. Moreover, we assess the impact of five popular imputation techniques, including scImpute, SAVER, Autoencoder (AE), Variational Autoencoder (VAE), and Generative Adversarial Network (GAN) on recovering biologically relevant co-expression patterns. Our results demonstrate that imputation models not only markedly enhance correlation between each promoter-reporter and native gene pair but also increase the number of cells co-expressing both genes. Imputation also improved transcription factor target gene correlations and revealed stronger associations among genes within the same protein complex. This work highlights the utility of promoter-reporter systems for benchmarking computational methods and underscores the potential of deep learning-based imputation to improve the biologically relevant signals of scRNA-seq data.

## Backgrounds

Single-cell RNA sequencing (scRNA-seq) improves upon traditional bulk RNA-seq by revealing transcriptomic differences between individual cells, enabling the construction of gene regulatory networks at the single-cell level^1–3^. Analyzing correlation patterns in single-cell gene expression data is fundamental for understanding co-expression networks, which help identify functional modules^4^. Constructing gene co-expression networks requires computational methods to calculate the correlation between the expression of different genes^5^. In the past two decades, various methods have been developed and applied to data from microarray and bulk RNA-seq, including Pearson’s linear correlation, biweight midcorrelation (bicor), Spearman’s rank correlation, Kendall’s Tau correlation, and mutual information^5–7^. However, these correlation measures have not yet been extensively applied or validated for single-cell RNA-seq data, where the inherent sparsity and cell-to-cell heterogeneity pose unique analytical challenges. To address these challenges, some specialized methods have been developed. For example, CS-CORE estimates gene-gene correlations within individual cell types, rather than across the entire dataset, allowing for more biologically meaningful co-expression inference by accounting for cell-type-specific expression patterns^8^. hdWGCNA extends the classical weighted gene co-expression network analysis (WGCNA) framework to handle the high dimensionality and sparsity of single-cell data, enabling the identification of co-expression modules at single-cell resolution^9^. Despite advancements in correlation-based methods, a systematic comparison of different correlation measures in single-cell RNA-seq data is still lacking.

Moreover, standard correlation analyses can also be affected by zero inflation, where genes exhibit artificially low or absent expression due to technical dropout rather than true biological absence. To mitigate this issue, pseudo-bulk analysis^10^, which aggregates gene expression across cell clusters, has been employed to reduce sparsity and enhance signal robustness. In addition, a variety of imputation methods have emerged as powerful tools to recover missing expression values and improve downstream analyses. Among these, scImpute and SAVER are widely used statistical approaches specifically developed for imputing dropout values in scRNA-seq data^11,12^. These methods have been ranked the best methods for single cell imputation in a recent benchmarking comparison^13^. The scImpute method estimates the true expression level of each gene in a cell by borrowing information from similar cells^11^, while SAVER uses a Bayesian framework to recover gene expression by leveraging prior information and smoothing across genes and cells^12^. Although both methods have demonstrated improvements in data quality, they are computationally intensive and often require substantial runtime, particularly for large single-cell datasets^14–17^. To address these limitations, deep learning-based methods, including Autoencoders (AE), Variational Autoencoders (VAE), and Generative Adversarial Networks (GAN) have been proposed; these methods offer computational efficiency and scalability while learning the underlying structure of the data to recover missing expression values and improve downstream analyses^18–21^. While these methods are promising, their relative performance in recovering biologically meaningful gene co-expression relationships has not been fully tested.

To address this gap, we developed scCoBench, a benchmarking study evaluating gene-gene correlation metrics and imputation methods using a biologically grounded reference system. Specifically, we generated four novel scRNA-seq datasets in *Arabidopsis thaliana* to investigate the relationship between transcript levels of promoter-reporter genes and their native counterparts. Green or Yellow Fluorescent Protein (GFP or YFP) reporter systems have long been used in plant biology research to study spatial and temporal gene expression patterns^22^. Because these reporters are driven by native promoters, their expression patterns closely mirror those of the corresponding native genes^23,24^. This makes promoter-reporter pairs a gold standard for benchmarking, as they serve as independent yet biologically linked proxies for native gene expression^23,24^. By using promoter-GFP or -YFP as internal control, they enable the precise evaluation of gene expression correlation measures, particularly in single-cell studies, where ground truth for gene-gene co-expression is often unavailable. We used the Arabidopsis root as our model system because of the rich resource of promoter-reporter lines, the wealthy of genomic knowledge for the gene expression patterns from many cell-type specific and single cell studies, the ease of observing the gene expression using confocal imaging, and the method of using cell sorting to validate our computational predictions^23–29^. To our knowledge, such internal controls have not been used in any species for benchmarking single-cell co-expression before. Leveraging this system as a case study, we assessed the performance of various gene-gene correlation measures in capturing relationships between promoter-reporter genes and their corresponding native transcripts. This dataset provides a valuable resource for benchmarking analytical methods and highlights key considerations for selecting computational tools for constructing gene co-expression networks in single-cell transcriptomic studies.

## Results

### Pipeline for quantifying and validating gene-gene relationships in single-cell RNA-seq data

To investigate gene relationships using scRNA-seq data, we established scCoBench, a multi-step analysis pipeline (Figure 1). The workflow begins with a raw gene expression matrix and proceeds through multiple analytical steps. Gene-gene relationships were first quantified using various similarity metrics, including six correlation measures (e.g., Pearson, Biweight midcorrelation, Spearman, Kendall, CS-CORE, p-transform), three distance metrics (e.g., Euclidean, Manhattan, Chebyshev), and a mutual information method. These calculations were applied across different subsets of data (i.e. all cells, expressing at least one, or both genes in a pair) and using different normalization strategies (i.e. including raw counts, log-normalized, or scaled data). In addition to calculating the co-expression using data from each single cell, to improve the robustness of the analysis, gene expression values were aggregated across cell types to create pseudo-bulk profiles that reduce cell-level noise. Moreover, imputation methods were employed to recover dropout values and enhance data quality. The impact of imputation was then assessed by comparing correlations between transcription factor (TF) target genes and genes in the same protein complexes before and after imputation, to evaluate whether biologically meaningful regulatory signals are captured. Finally, predicted gene-gene relationships were validated experimentally, such as through PCR-based assays, to confirm that highly co-expressed genes identified by scCoBench can be detected in the same cell type.

**Figure 1:**
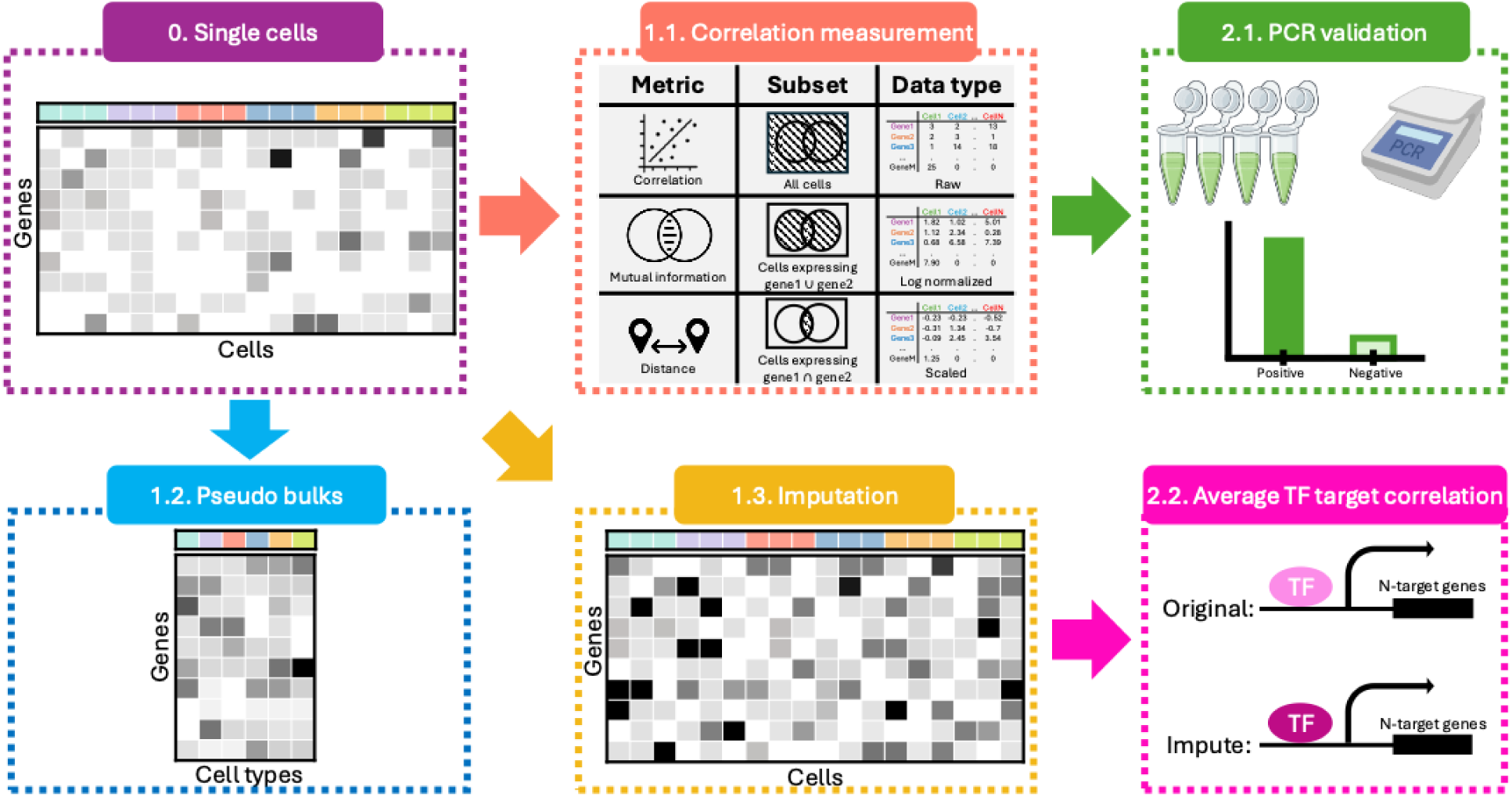
Pipeline for analyzing gene relationships using single-cell RNA-seq data. • <u>Step 0</u>: a raw single-cell gene expression matrix. • <u>Step 1.1</u>: Gene-gene relationships are quantified using various metrics such as correlation (e.g., Pearson, Biweight midcorrelation, Spearman, Kendall, CS-CORE, p-transform), mutual information, or distance metrics (e.g., Euclidean, Manhattan, Chebyshev). These analyses are performed across different data subsets (e.g., all cells, cells expressing either or both genes in a pair) and data normalization types (raw count, log normalization, or scaled data). • <u>Step 1.2</u>: Gene expressions are aggregated by cell types to create pseudo-bulk profiles before downstream analysis. • <u>Step 1.3</u>: Gene expression matrices are imputed to recover dropout values and enhance signal quality. • <u>Step 2.1</u>: Predicted gene co-expressions are validated experimentally using PCR assays. • <u>Step 2.2</u>: The impact of imputation is assessed by comparing transcription factor (TF)-target gene correlations before and after imputation.

To demonstrate the utility of our pipeline, we conducted a comprehensive analysis comparing the expression patterns of four Arabidopsis root promoter-reporter lines - *pWER:GFP*, *pSCR:GFP*, *pCORTEX:GFP*, and the PET111 enhancer trap - with their corresponding native genes using single-cell RNA-seq data (Supplementary Figure 1A-D). This comparison enabled us to assess how accurately promoter-reporter constructs reflect native gene expression at single-cell resolution. *pWER:GFP* marks the epidermis and lateral root cap, consistent with the expression of (WER)^25^. *pSCR:GFP* labels the endodermis and quiescent center, aligning with the expression of SCARECROW (SCR)^26,27^. *pCORTEX:GFP* specifically labels the cortex layer and corresponds to AT1G09750, a known marker gene for cortical cells which is confirmed by single-cell RNA-seq data^28^. PET111 is an enhancer trap line that drives YFP expression in the columella; while it does not have a defined native gene, its likely regulatory target was inferred by aligning its spatial expression pattern with columella specific gene expression profiles in the single cell dataset^29^. The FeaturePlot revealed strong concordance between the expression patterns of native genes, and their respective promoter-reporter constructs across all samples (Figure 2A, Supplementary Figure 2A-C), supporting the use of these datasets as robust standards for benchmarking correlation methods.

**Figure 2:**
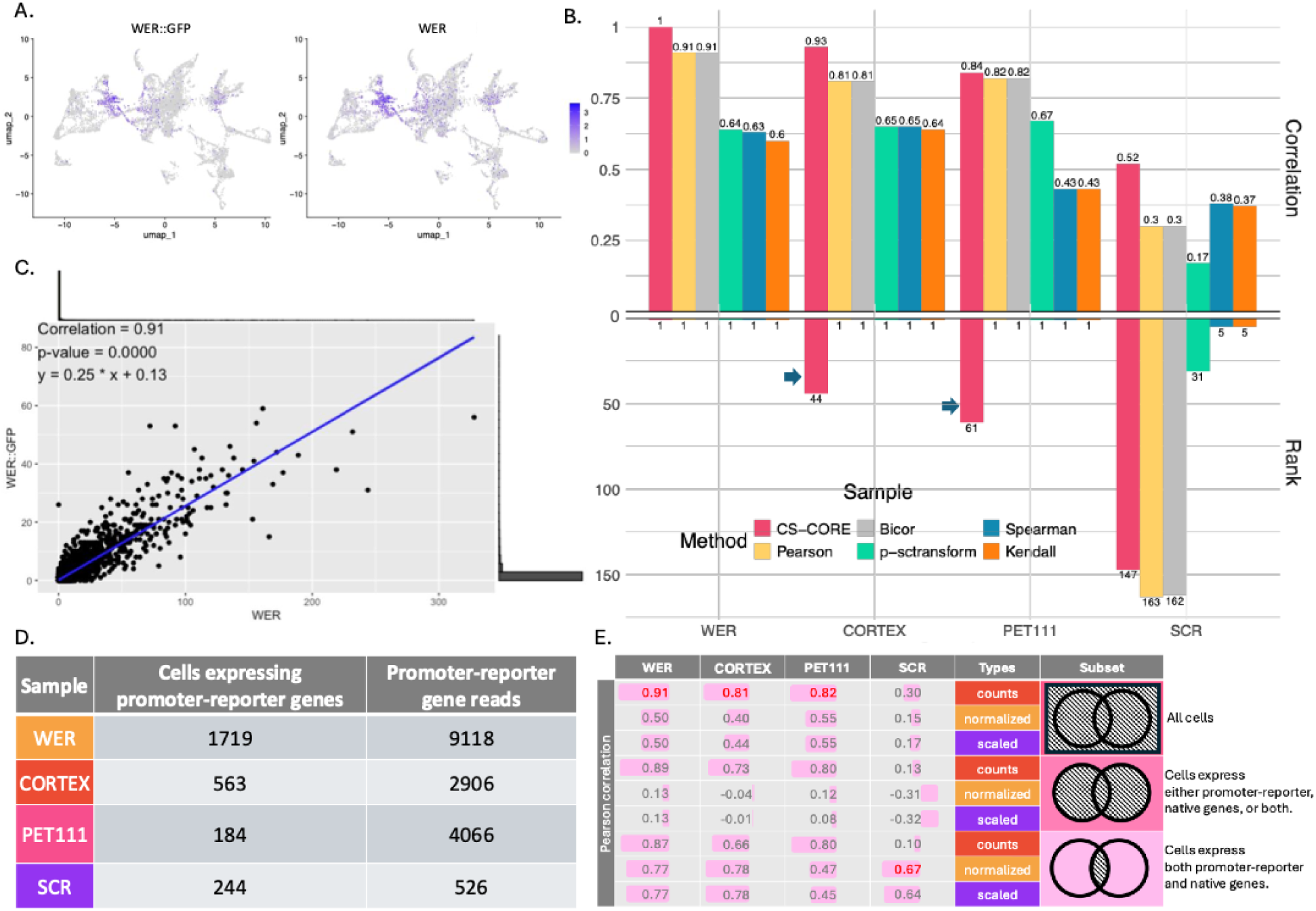
Comparative analysis of promoter-reporter and native gene expression across different methods and samples. A. Feature plot showing the gene expression patterns of *pWER:GFP* and the WER gene. B. The top bar chart shows the correlation between promoter-reporter and native genes for each sample using six correlation methods. The bottom bar chart displays the rank correlation of promoter-reporter and native genes in each sample. C. Scatter plot and the Pearson correlation between *pWER:GFP* and the WER gene. D. The number of cells expressing promoter-reporter genes and the total number of reads in each sample. E. The correlation between promoter-reporter and native genes across different data types (counts, normalized, and scaled) and data subsets (all cells, cells expressing either promoter-reporter or native genes, or both, and cells expressing both promoter-reporter and native genes).

### Benchmarking correlation methods to evaluate promoter-reporter and native gene relationships in single-cell RNA-seq data

To quantify the relationship between promoter-reporter and native gene expression, a comprehensive correlation analysis was conducted using six correlation methods: CS-CORE, Pearson, Bicor (Biweight midcorrelation), p-sctransform, Spearman, and Kendall (Figure 2B). We also employed mutual information and three matrix distance methods (Supplementary Figure 3A, B). Among these correlation methods, CS-CORE initially demonstrated the highest correlation between promoter-reporter and native genes across all four samples: WER (1), CORTEX (0.93), PET111 (0.84), and SCR (0.52). However, further analysis revealed limitations in the CS-CORE method when dealing with lowly-expressed genes or genes expressed only in specific regions, as observed in the CORTEX and PET111 samples. While other methods consistently ranked the promoter-reporter genes as having the highest correlation with their corresponding native genes (Figure 2b, Supplementary Figure 3A, B, C), CS-CORE ranked the correlation between *pCORTEX:GFP* and its native gene (AT1G09750) at 44th, and *pPET111:YFP*’s correlation with its corresponding gene (AT2G28870) at 61st (blue arrow, Figure 2B). Despite this lower ranking, this result from CS-CORE is expected because this method is designed for calculating co-expression at the cell-type-specific level, which is different from other methods which are calculating co-expression across all cells.

Among the correlation-based methods, Pearson and Bicor methods generated the highest co-expression for three out of four gene pairs. Spearman and Kendall methods generated higher correlations and rankings between SCR reporter and SCR gene than Pearson and Bicor methods. Regardless of the methods used, the correlations of SCR reporter and SCR gene are always lower than other gene-reporter pairs.

Although WGCNA is one of the most widely used methods for identifying co-expression modules, interestingly, hdWGCNA did not capture the strong relationship between promoter-reporter and native genes. While the reporter and native genes in each sample were assigned to the same co-expression module, indicating shared network structure, their TOM values (a topological measurement for gene-gene similarity used by WGCNA/hdWGCNA) were relatively low. More importantly, the reporter genes did not rank among the top co-expressed partners of their corresponding native genes based on TOM values (Supplementary Figure 4A, B, C, D, E). However, hdWGCNA did successfully group the reporter and the native genes in the same module, which is also the expected behavior of this method (Supplementary Figure 4A, B, C, D, E).

Unlike all other methods tested, mutual information used an information-theoretical approach to determine gene-gene relationships. Our analysis showed that mutual information also confirmed the strong relationship between promoter-reporter and native genes (Supplementary Figure 5A), aligning with the results from other correlation methods. We also explored distance-based methods, including Euclidean, Manhattan, and Chebyshev distances, to assess the similarity between gene expression profiles. Interestingly, these distance metrics did not consistently identify the promoter-reporter gene as the most similar to its corresponding native gene (Supplementary Figure 5B). This discrepancy highlights the potential limitations of using distance-based approaches alone for assessing gene expression similarities in this context. These findings underscore the importance of employing multiple analytical approaches to ensure robust characterization of gene expression relationships in single-cell studies.

To further visualize these relationships, we generated scatter plots for each gene pair. The plot for *WER* and *pWER:GFP* revealed a strong positive Pearson correlation of 0.91 (Figure 2C). Accompanying histograms on the upper and right sides of the scatter plot illustrated the distribution of cells expressing the native and promoter-reporter genes. These histograms uncovered a significant challenge in our study: a large proportion of cells did not express either the native or the promoter-reporter genes across all samples. This issue was particularly pronounced in the *pSCR:GFP* sample, which exhibited notably low expression levels with 526 reads in more than 660 million reads from this sample (Figure 2D, Supplementary Figure 1E). This finding potentially explains the markedly lower correlation observed in the *pSCR:GFP* sample compared to the others.

To explore the impact of non-expressing cells on correlation measurements, we measured correlations in various subsets of the data: (1) all cells, (2) cells expressing either the promoter-reporter or native gene or both, and (3) cells expressing both genes (Figure 2E). Additionally, three data types were compared: raw read counts, normalized data, and scaled data (Fig 2E). Surprisingly, native and promoter-reporter genes showed the highest correlation (0.91, 0.81, and 0.82 for WER, CORTEX, and PET111, respectively) in the raw count data of all cells (Fig 2E). The *pSCR:GFP* and SCR gene pair, however, exhibited the highest correlation in the normalized data of cells expressing both genes (Fig 2E). Spearman and Kendall show the highest correlation between promoter and reporter genes in the subset of cells expressing both genes (Supplementary Figure 6A, B). This analysis highlighted that cells with no expression or very low expression could significantly affect correlation measurements. An earlier statistical study of correlation for zero-clustered data in medical research^30^ demonstrated that the Pearson’s method on count data can better estimate the true correlation than rank-based approaches, which partly aligns with our empirical observation in this study.

To further assess the robustness of our findings, we simulated scRNA-seq data using scDesign2^31^, a model-based simulator designed to preserve gene-gene correlation structures. In all four samples, scDesign2 successfully recapitulated the expression trends observed in the original data. Notably, Pearson correlation showed stronger within-cluster co-expression compared to Spearman n the original data. This trend was consistent in the simulated data, where Pearson also captured higher within-cluster correlations. (Supplementary Figure 7A, B, C, D). These simulation results and the results from our benchmarking experiments (Figure 2) demonstrate that Pearson correlation provides a robust metric for capturing gene-gene relationships, even in the presence of data sparsity or variability across cell types.

### Improving correlation robustness in single-cell data through pseudo-bulk analysis

To mitigate the impact of zero-inflation (the observation of excessive number of zeros) in single cell data, a pseudo-bulk analysis was implemented, averaging gene expression within clusters (Fig 3A, Supplementary Figure 8A). Pseudo-bulk approach has been recommended for testing differential gene expression analysis^10^, however, the effect of this approach on co-expression analysis has not been tested before. Two clustering approaches were compared: the default clusters using Seurat pipeline (Fig 3C) and cell type clusters labeled by the ortho-marker-groups (OMGs) method^32^ (Figure 3B, D, Supplementary Figure 8B). Pearson correlation demonstrated consistently and statistically significant high correlations (above 0.8) across all samples and data types (count, normalized, and scaled) in both clustering approaches (top bar chart Figure 3E and F). In contrast, Spearman correlation showed variability in different genes: the correlation is high in the WER sample, moderate in PET111 and SCR, and low in CORTEX for default clustering. With cell type clustering, Spearman correlation improved but still showed low correlation in the CORTEX sample (bottom bar chart Figure 3E and F). Kendall correlation (another rank-based method) aligned with Spearman results (Supplementary Figure 9). The p-sctransform method showed high correlations for all samples, validated by statistical tests (Supplementary Figure 9). CSCORE exhibited high correlations across all samples but lacked statistical significance (Supplementary Figure 9). Notably, cell type clusters generally yield higher correlations compared to default clusters, suggesting that biologically informed clustering results in stronger correlations. Despite the improvements offered by pseudo-bulk analysis, we recognized that the relatively low number of clusters could pose a challenge for achieving statistical significance (for example, see sample CORTEX, Figure 3E and F).

**Figure 3.**
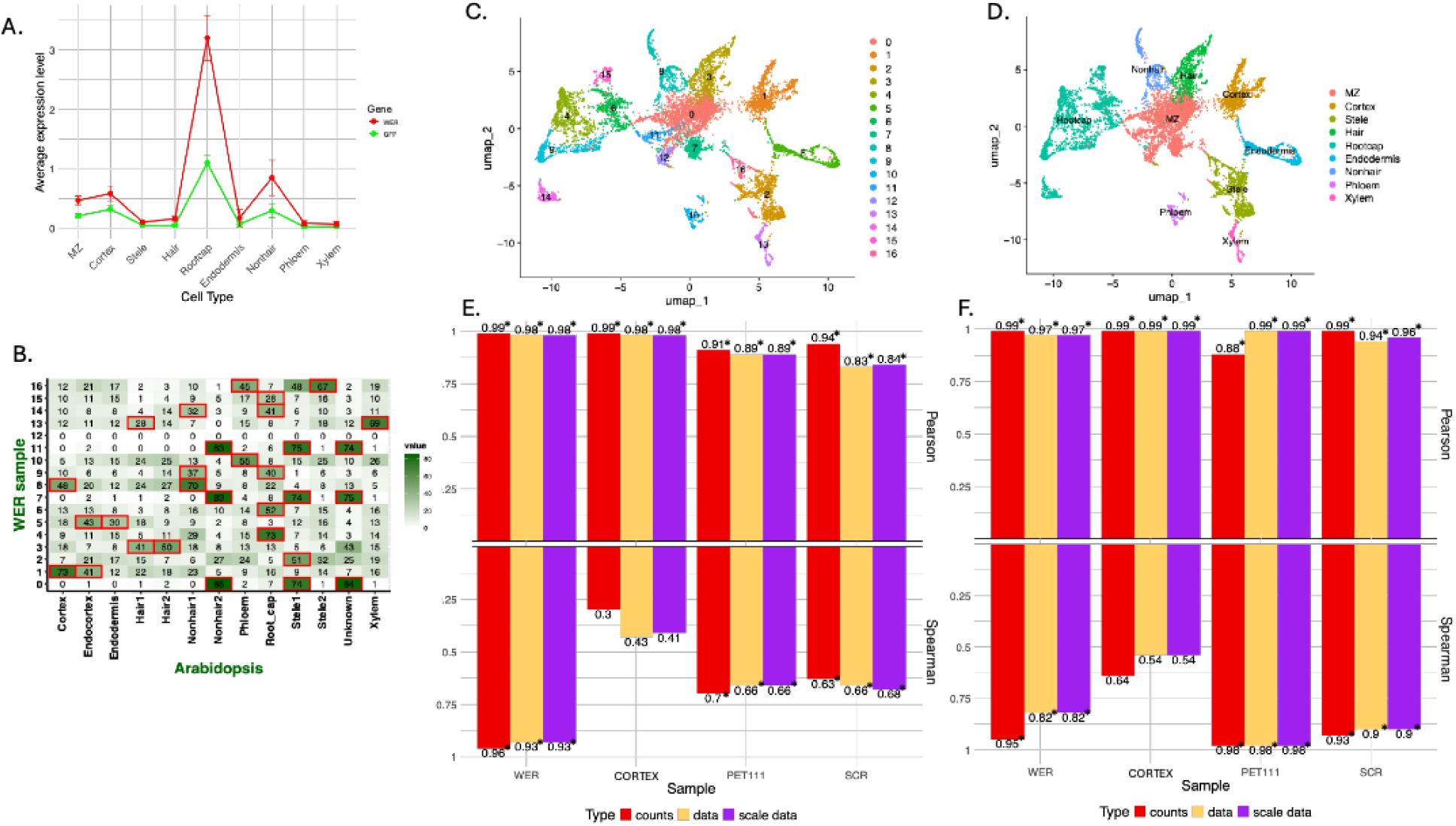
Promoter-reporter and native gene expression in pseudo-bulk analysis. A. The average gene expression patterns of *pWER:GFP* and the WER gene across different cell types in the sample. B. Cell type prediction for each cluster in the WER sample using the OMG method. The red boxes highlight statistically significant overlapping of marker genes with a reference single cell map, which is used to name the clusters for pseudo-bulk analysis. C. The default UMAP clusters generated by the Seurat pipeline. D. The cell type clusters as predicted by the OMG method. E. Correlation between the average gene expression of promoter-reporter and native genes in the default clusters, using Pearson (top) and Spearman (bottom) methods. F. Correlation between the average gene expression of promoter-reporter and native genes in the cell type clusters, using Pearson (top) and Spearman (bottom) methods.

### Mitigating dropout effects with deep learning and statistical imputation methods in single-cell RNA-seq data

Another method to address the issue of zero-inflated data is data imputation. To this end, we applied five different imputation methods, including scImpute, SAVER, Autoencoder (AE), Variational Autoencoder (VAE), and Generative Adversarial Network (GAN). Our initial analysis without imputation revealed that only a subset of cells co-expressed both the native gene and its corresponding promoter-reporter. For example, between WER and pWER:GFP, there is an approximately 45% overlap, contrary to expectations of a higher concordance (Figure 4A). Histograms further indicated that a large proportion of cells had low expression levels, often only one read per cell (highlighted by red arrows), suggesting that technical dropout might missed true biological expression of the target genes (Figure 4A).

**Figure 4.**
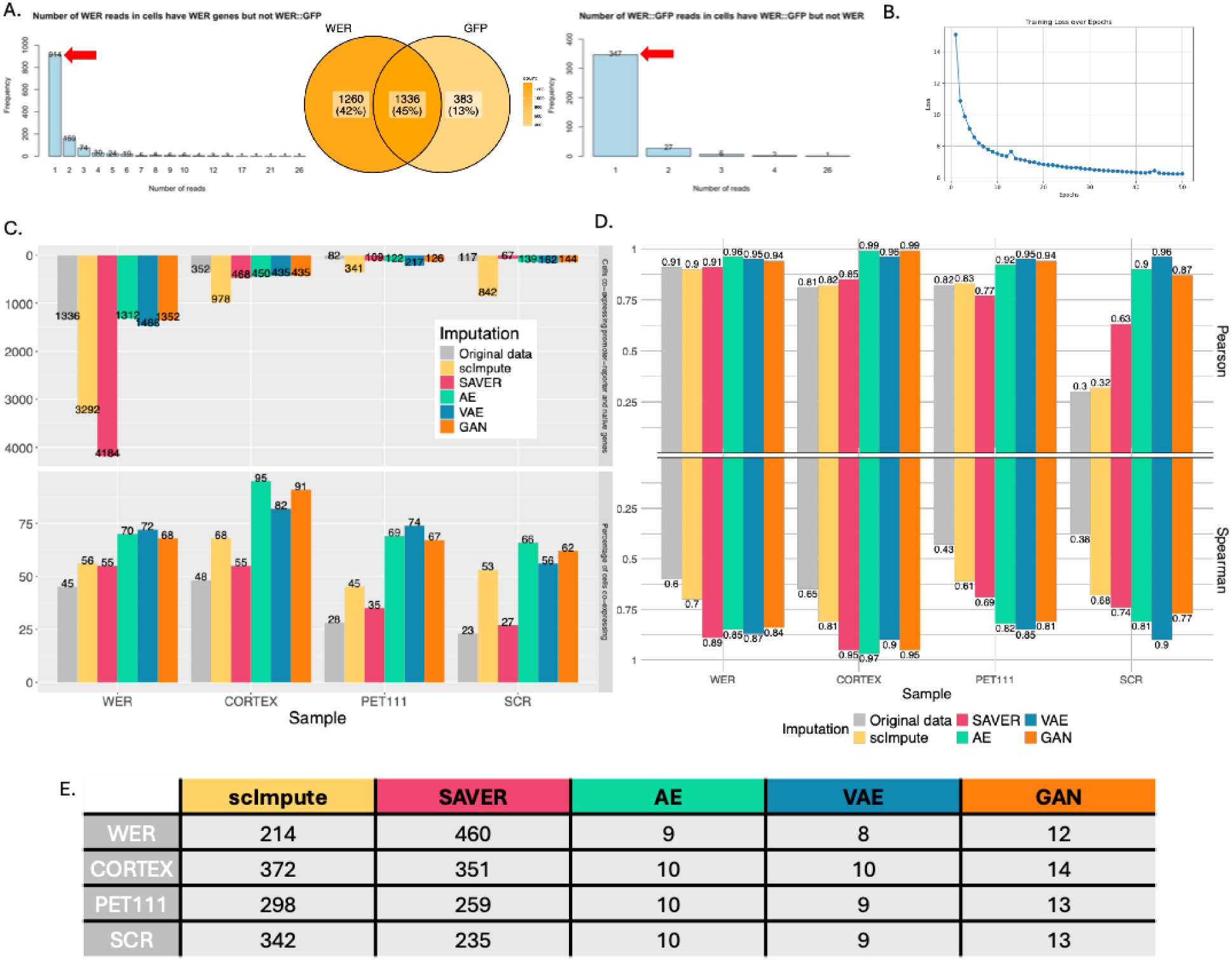
Evaluating imputation methods for promoter-reporter and native gene expression. A. The Venn diagram shows the number of cells expressing either WER, *pWER:GFP*, or both. The histogram on the left displays the distribution of WER reads in cells expressing WER but not *pWER:GFP*, while the histogram on the right shows the distribution of *pWER:GFP* reads in cells expressing *pWER:GFP* but not WER. B. Reconstruction loss of VAE in the WER sample. C. The top bar chart shows the number of cells expressing both the promoter-reporter and native gene in the original data and in the imputed data using AE and VAE methods across all samples. The bottom bar chart shows the percentage of cells expressing both the promoter-reporter and native gene in the original and imputed data by AE and VAE methods across all samples. D. The correlation between promoter-reporter and native genes is displayed for original and imputed data using AE and VAE across all samples, with Pearson correlation on the top and Spearman correlation on the bottom. E. Running time (in minutes) across imputation methods (scImpute, SAVER, AE, VAE, GAN)

We tested scImpute and SAVER, two popular methods for imputation that were ranked as top performers in a recent benchmarking study. Unlike scImpute and SAVER, AE, VAE, and GAN are deep learning-based approaches that can be easily implemented using the Pytorch framework, and we trained these models from scratch in this study. Other established VAE or GAN methods, such as scVI^33^ or cscGAN^20^, did not provide the flexibility of imputing all genes in the data in their implementations and were not compared in this study. To assess the performance of these models, we monitored reconstruction loss during training. As an example, the VAE model demonstrated stable convergence across epochs (Figure 4B), supporting its effectiveness in recovering expression patterns through simple neural network models.

Following imputation, we observed a substantial increase in the number of cells expressing both the native and promoter-reporter genes, particularly with the scImpute and SAVER methods (Figure 4C, top panel, Supplementary Figure 10). These improvements were consistent across all four samples (WER, CORTEX, PET111, and SCR), with the most pronounced effect observed in the SCR sample by scImpute. Similarly, the percentage of cells co-expressing both gene types also increased in the imputed data (Figure 4C bottom panel, Supplementary Figure 10), particularly higher in VAE, AE, and GAN models. Importantly, unlike scImpute and SAVER, VAE/AE/GAN models increased the overlapping percentages, without substantially increasing the number of cells expressing both reporter and native genes.

These imputation methods also resulted in higher correlation between promoter-reporter and native gene expression. Both Pearson and Spearman correlations increased markedly after imputation, with AE and VAE achieving the most consistent enhancements across samples (Figure 4D). The largest improvement was again observed in the SCR sample, where the Pearson correlation increased from 0.30 in the original data to 0.96 after VAE imputation. Finally, we also evaluated the computational efficiency of each method, finding that deep learning model training from scratch like AE, VAE, and GAN required significantly less runtime compared to scImpute and SAVER (Figure 4E). These results highlight the effectiveness of deep learning imputation approaches in mitigating dropout effects and improving the accuracy of gene-gene relationship analysis in single-cell transcriptomic data.

### Validating biological relevance of imputed data

To assess the biological relevance of our imputed data, we calculated correlations between transcription factors (TFs) and their target genes. Only VAE results were shown here for simplicity. For each TF, we calculated pairwise correlations among its known target genes (based on a published DAP-seq dataset)^34^ in both the original and imputed datasets. The average of the pairwise correlation between these target genes was higher in the imputed data compared to the original data (Figure 5A). We also observed a positive relationship between target gene expression levels and their correlations in both the original and imputed datasets (Figure 5B). Interestingly, while the imputation process did not artificially inflate gene expression levels (Figure 5C), it did lead to significantly improved correlations among target genes (Figure 5D, p<0.01). This finding indicates that the imputation methods were able to recover biologically meaningful signal from the noisy single-cell data without introducing spurious high expression values. Our findings provide strong evidence for the effectiveness of imputation methods in recovering biologically relevant information from scRNA-seq data.

**Figure 5.**
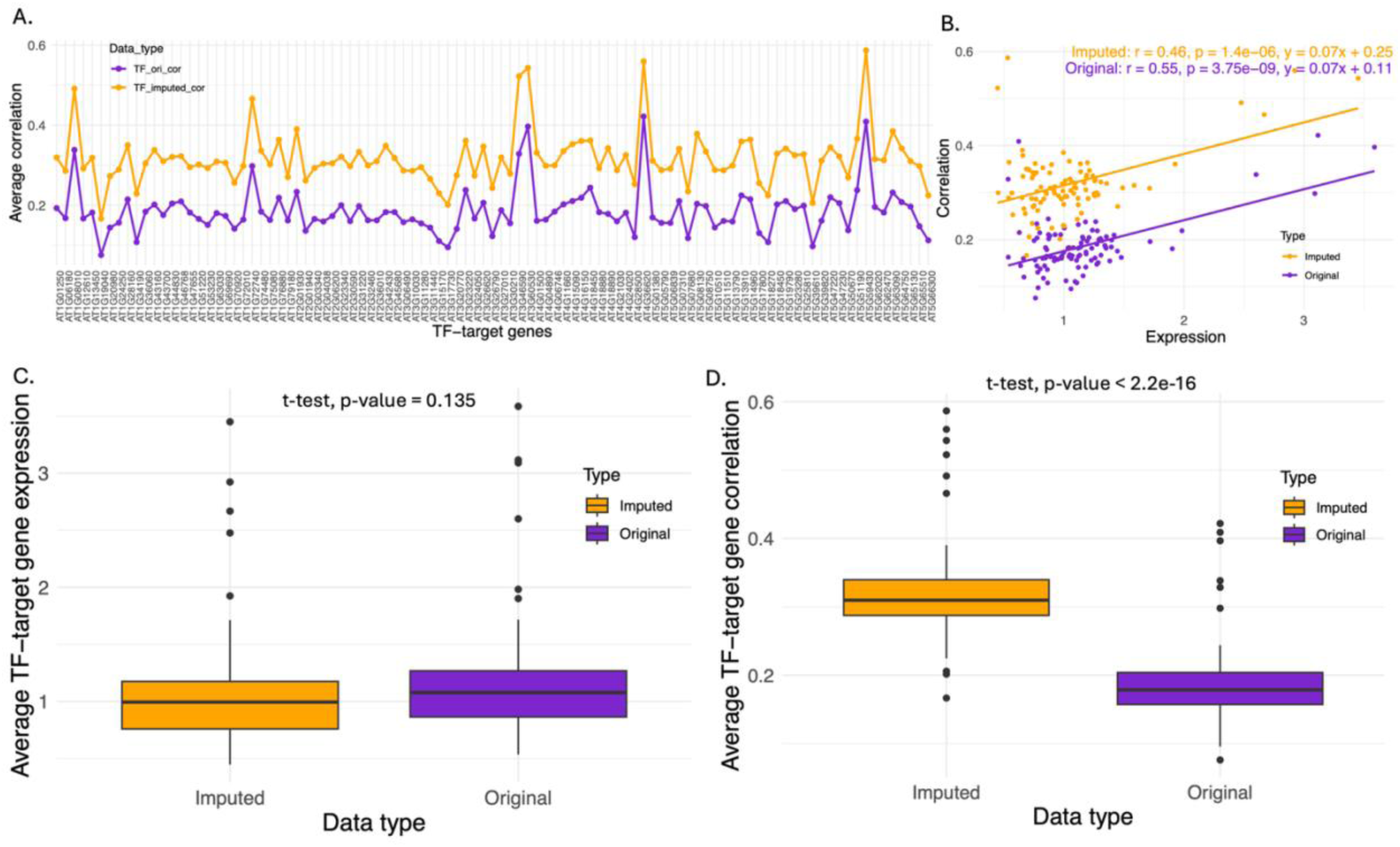
Impact of imputation on transcription factor target gene correlation. A. The line plot displays the average correlation of target genes for each TF in the original data (purple) and imputed data (yellow). B. Scatter plots with fitted lines illustrate the relationship between gene expression and correlation for each TF, represented by dots. Purple indicates the original data, yellow the imputed data. The correlation strength, p-value, and best-fit line are shown for each dataset. C. The boxplot compares the average target gene expression for each TF. A t-test was performed, but no significant difference was found between imputed and original data expression. D. The boxplot compares the average target gene correlation for each TF. A t-test revealed significantly higher correlation in the imputed data compared to the original data.

To further evaluate the impact of imputation in a different application context, we analyzed correlations between nucleoporin genes and other genes associated with the nuclear pore complexes (NPC), including genes involved in transcriptional regulation, mRNA splicing and processing, translational regulation, and a set of randomly selected genomic genes^35^. This recently published dataset was chosen because the approach utilizes proximity labeling, which provides improved assessment of true protein complex associations as compared to traditional proteomics approaches. Using imputed data from VAE, both Pearson and Spearman correlation analyses revealed stronger correlations than the original analysis (Figure 6), further supporting the biological validity and utility of imputation in single-cell transcriptomic analyses.

**Figure 6.**
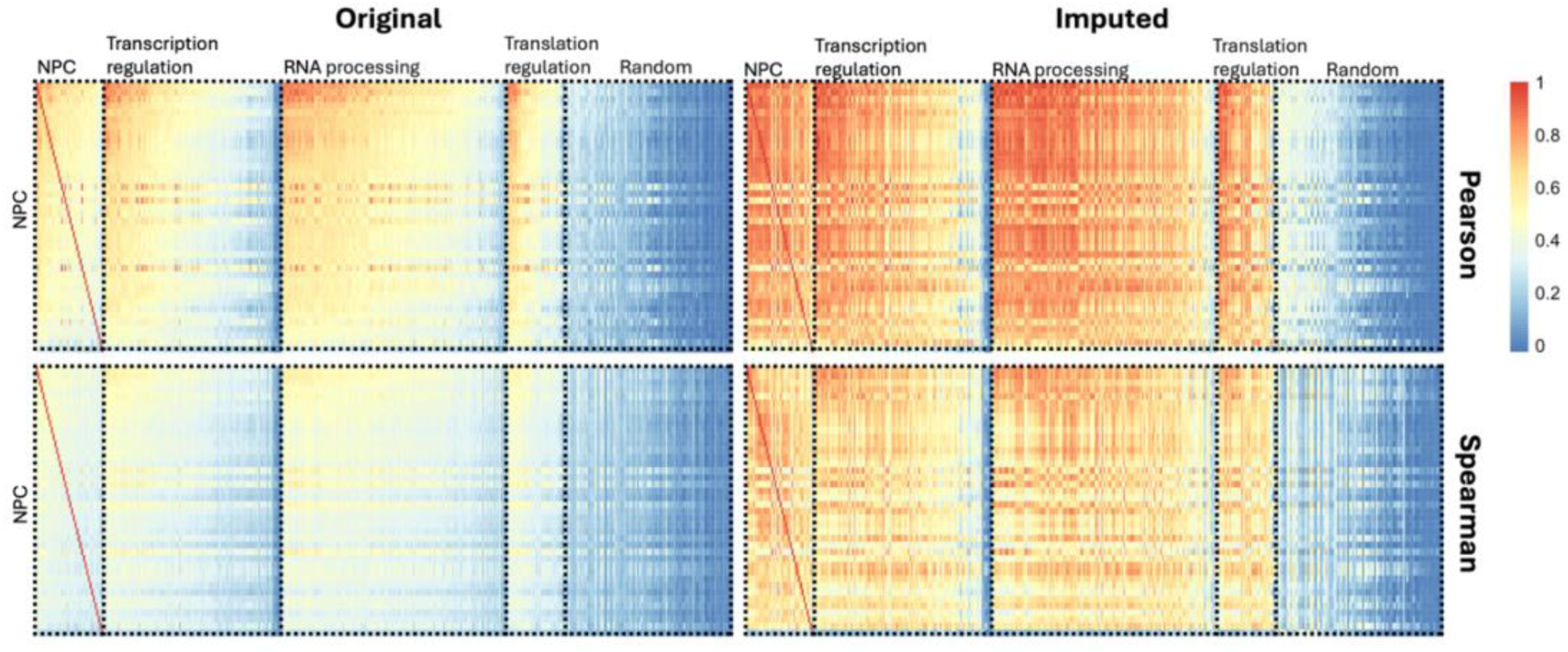
Correlation analysis between nucleoporin genes and associated functional components in original and imputed data. Pearson and Spearman correlations between nucleoporin genes (NPC) and other NPC-associated components, including transcription regulation, mRNA splicing and processing, translation regulation, and randomly selected genes from the genome, in both original and imputed datasets. The top heatmaps represent correlations computed using Pearson correlation, while the bottom heatmaps represent correlations computed using Spearman correlation. NPC genes showed high correlation with biologically relevant genes in transcription regulation, translation regulation and RNA processing and low correlation with randomly selected background genes.

### Experimental validation of correlation findings

Using our correlation-based approach, we identified AT1G70370 and SMR10 (AT2G28870) as genes highly co-expressed with the reporter constructs *pCORTEX:GFP* and *pPET111:YFP*, respectively. The pCORTEX:GFP is from a known gene promoter, thus we select the highest correlated gene (AT1G70370) as the target for validation. The *pPET111:YFP* is an enhancer trap line with an unknown insertion site. We performed whole-genome resequencing, and the result did not reveal any nearby gene closely linked to the reporter insertion site. To validate these predictions, qRT-PCR were performed and the results confirmed that both genes are significantly enriched in reporter-labeled (GFP+ or YFP+) cells, with minimal expression in non-labeled populations. The strong expression of SMR10 in the YFP-sorted cells and AT1G70370 in the GFP-sorted cells supports that these genes are co-expressed in the same cell type as predicted. These results further validate the specificity of the reporter lines and support the biological relevance of the co-expression relationships inferred from our single-cell analysis (Figure 7).

**Figure 7.**
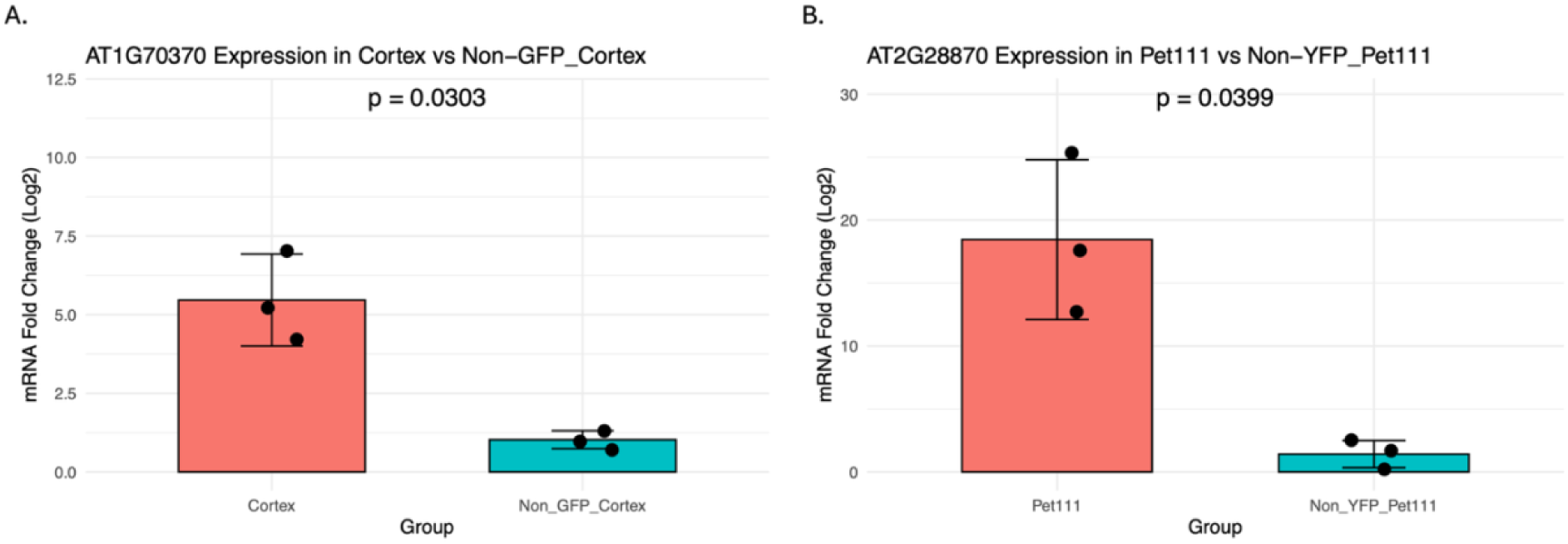
Quantitative PCR (qPCR) validation of top correlated gene expression in GFP/YFP-labeled and non-labeled cell populations from Cortex and PET111 reporter lines. A, B: Bar plots show the expression levels of the genes most highly correlated with the reporter constructs in the Cortex and PET111 lines, measured by qRT-PCR in GFP/YFP-positive versus non-labeled cells. Each dot represents a biological replicate; error bars indicate standard deviation. For both lines, the genes selected for validation were identified through correlation analysis of single-cell RNA-seq data as the top co-expressed genes with the reporter signal.

## Discussion

Understanding gene-gene relationships at single-cell resolution is crucial for decoding complex regulatory networks and biological processes. In this study, we presented scCoBench, a comprehensive pipeline for quantifying and validating gene-gene relationships using single-cell RNA-seq (scRNA-seq) data. By benchmarking multiple correlation and distance-based similarity metrics across different sampling strategies, pseudo-bulk aggregation, and a range of imputation techniques including deep learning models, we provided a robust framework for assessing the quality and biological relevance of gene expression relationships.

Our benchmarking analysis using four Arabidopsis root promoter-reporter lines, pWER:GFP, pSCR:GFP, pCORTEX:GFP, and the pPET111:YFP enhancer trap, demonstrated strong concordance between reporter constructs and their corresponding native genes. This result affirmed the utility of promoter-reporter systems as biologically meaningful standards for validating computational approaches. We focused on root tissues due to the availability of well-characterized cell type-specific reporters, the established and highly ordered cellular architecture of the Arabidopsis root, and the abundance of publicly available root scRNA-seq datasets^23^. Moreover, root tissues exhibit more stereotyped spatial expression patterns than other plant organs and allow earlier, more precise collection of developmental stages^36^. These features make the Arabidopsis root an ideal model for systematically evaluating gene-gene relationships in a controlled and biologically interpretable context.

Among the correlation methods tested, Pearson correlation consistently ranked the native-promoter gene pairings highly and remained robust across raw, normalized, and scaled data. We observed that zero-inflation and sparse co-expression across cells significantly impacted correlation strength. To further mitigate the impact of sparsity, we applied pseudo-bulk profiling, which substantially improved correlation between promoter-reporter and native genes. Notably, biologically informed cell type clustering yielded higher and more consistent results than unsupervised clustering, suggesting the importance of incorporating cell identity information in single-cell co-expression analysis.

Beyond pseudo-bulk strategies, we evaluated several imputation methods to recover biologically meaningful expression affected by dropout events. Deep learning-based models, particularly autoencoders (AE) and variational autoencoders (VAE), showed superior performance in restoring gene expression patterns, increasing the proportion of co-expressing cells, and significantly enhancing correlation metrics between native and promoter-reporter genes. Our evaluation of imputation quality using transcription factor (TF) target gene co-expression and nucleoporin-associated gene correlations further confirmed that these methods recover biologically relevant signals without introducing excessive bias.

To experimentally validate the computational framework, we focused on PET111, an enhancer trap line without a defined native gene. Using our correlation-based pipeline, we identified SMR10 (AT2G28870) as the most highly co-expressed candidate gene. This prediction was supported by qRT-PCR, which demonstrated strong expression of SMR10 in PET111-labeled cells and negligible expression in negative controls, confirming the specificity of our approach.

While this study introduces a biologically grounded benchmark for evaluating gene-gene correlation metrics and imputation strategies, several limitations should be considered. One limitation is the inconsistency in gene ranking across samples, even when using the same correlation method. This variation may arise from both technical noise and biological batch effects, highlighting the sensitivity of rank-based assessments to sample-specific factors. As a result, the top co-expressed partners of a gene may differ between datasets, complicating efforts to identify universally conserved gene relationships. This challenge may be addressed by integrating data across samples using batch correction methods that preserve biological variation while minimizing technical differences^37^ (Supplementary Figure 11).

Another consideration is that the performance of correlation methods in this study was not assessed under dynamic or perturbed conditions, such as time-series data or multi-condition experiments, where gene-gene relationships may shift in response to environmental or genetic changes. Although pseudo-bulk strategies enhanced signal stability, they also reduced resolution and statistical power due to the limited number of clusters in certain samples. This trade-off may obscure subtle or rare cell type specific co-expression patterns.

Imputation methods also come with caveats. While they helped recover gene expression and improved correlation scores, they rely on certain statistical assumptions about the data. These assumptions can unintentionally distort the true expression landscape by artificially increasing similarity between genes or smoothing over genuine biological variability^38^. Therefore, imputation may introduce biases that affect downstream analyses. A potential solution is selective imputation, where dropout recovery is applied only to low-expression values while preserving the natural variance of moderately or highly expressed genes^16^.

Our study was conducted exclusively in Arabidopsis thaliana using protoplast-based scRNA-seq data, and conclusions may not generalize to other plant species or animal systems. Additional studies in diverse biological contexts are needed to evaluate the robustness and transferability of the methods presented here. Lastly, experimental validation was limited to a subset of gene pairs. While qRT-PCR confirmed strong co-expression for selected cases, broader validation across more cell lineages could help improve the assessment the biological accuracy of predicted gene-gene relationships.

## Conclusion

The scCoBench pipeline provided a flexible framework for characterizing gene-gene relationships in scRNA-seq data, accounting for data sparsity, gene expression heterogeneity, and dropout effects. The integration of promoter-reporter benchmarks, multiple correlation measures, and deep learning-based imputation highlights the potential of computational tools to improve the resolution and interpretability of single-cell datasets. Our results highlighted the importance of choosing appropriate correlation methods, considering the impact of non-expressing cells, and employing advanced techniques like pseudo-bulk analysis and data imputation to overcome the challenges inherent in scRNA-seq data. The study demonstrated that these approaches can significantly enhance our ability to accurately measure and interpret gene-gene relationships in single-cell transcriptomics, opening new avenues for understanding gene regulation and expression dynamics at the cellular level.

## Methods

### 1. Plant materials and preparation of scRNA-seq samples

Transgenic lines of Arabidopsis thaliana used in this study were as previously described; *pWER:GFP^25^*, *pSCR:GFP^26,27^*, *pCORTEX:GFP^28^*, *pPET111:YFP^29^*. Sample preparation for scRNA-seq was performed as described in our previous publication^39^. Arabidopsis thaliana seeds were surface sterilized using a 30% (v/v) bleach, 0.1% (v/v) Triton X-100 solution for 10 min and incubated on Murashige and Skoog (MS) growth media covered with 100/47 mm mesh. The square dishes are oriented vertically and incubated under continuous light condition at 22°C for 4∼5 days after gemination (∼ 2-2.5cm long roots). At 4∼5 d after germination, the primary root tips were cut 1/3 from root tips and put them into a 35mm petri dish containing a 70um strainer and 4 mL enzyme solution (1.25% [w/v] Cellulase [“ONOZUKA” R-10, Yakult], 0.1%[w/v] Pectolyase [P-3026, Sigma-Aldrich], 0.4 M Mannitol, 20 mM MES [pH 5.7], 20 mM KCl, 10 mM CaCl2, 0.1% [w/v] bovine serum albumin). The sample was incubated on the orbital shaker at 85 rpm for 1 hour at 25°C. After incubation, the protoplast solution was filtered through a 70μm filter, and then through a 40μm filter twice in 50ml conical tubes. The filtered solution was centrifuged at 500g for 10 min at 22°C using a swinging bucket rotor and discard supernatant. The protoplasts were resuspended using 500μL Solution A (0.4 M Mannitol, 20 mM MES [pH 5.7], 20 mM KCl, 10 mM CaCl_2_, 0.1% [w/v] bovine serum albumin) and transferred to a 2ml round bottom tube. After centrifugation at 200g for 6 min, the pelleted protoplasts resuspended with 30-50 μL Solution A to achieve the desired cell concentration (∼1,000 protoplasts/μL) after cell counting.

### 2. Single-cell RNA-seq library construction and sequencing

The protoplasts were loaded into Chromium microfluidic chips (Next GEM Single Cell 3’ kit) and barcoded with a 10X Chromium Controller (10X Genomics). The mRNA from barcoded cells was then reverse-transcribed, and sequencing libraries were prepared using reagents from a Chromium Single Cell 3’ v3.1 reagent kit (10X Genomics) following the manufacturer’s guidelines. The cDNA and final library quality were confirmed using a Bioanalyzer (Agilent). Sequencing was performed with Illumina NovaSeq 6000 according to the manufacturer’s instructions.

### 3. Confocal Microscopy

Seedlings of Arabidopsis (4∼5 days after gemination) were stained with 10 µg/mL propidium iodide solution for 1 min and rinsed in the water to eliminate any remaining staining solution. Afterward, the stained roots were examined with a Leica SP5 laser scanning confocal microscope. The excitation wavelength was 488 nm for the detection of GFP signals and 561 nm for the propidium iodide.

### 4. Cell ranger mapping

Raw single-cell RNA-seq data were processed using “Cell Ranger v7.1.0” (10x Genomics). The “cellranger count” pipeline was used to align reads, generate gene expression matrices, and estimate the number of cells using the “--expect-cells” parameter. For each sample, sample-specific FASTQ files were provided via the “—fastqs” parameter, and the corresponding sample name was specified using the “—sample” parameter.

Before running “cellranger count”, a custom Arabidopsis thaliana transcriptome reference was generated using the “cellranger mkref” command. The reference was built using Ensembl Plants release 59, which included the GFF3 annotation file “Arabidopsis_thaliana.TAIR10.59.gff3.gz” and the protein FASTA file “Arabidopsis_thaliana.TAIR10.pep.all.fa.gz”. The GFF3 file was first converted to GTF format using “gffread” prior to reference construction. The resulting gene expression matrices were stored in the “outs/filtered_feature_bc_matrix.h5” file and used for downstream analyses in Seurat.

### 5. Data processing using Seurat

Raw scRNA-seq data for WER, SCR, PET111, and CORTEX samples were processed using “Seurat 5.1.0” in R. Gene expression matrices were loaded from 10x Genomics .h5 files using Read10X_h5(), and Seurat objects were created with CreateSeuratObject(). Quality control (QC) was performed by filtering cells with fewer than 500 genes or exceeding sample-specific thresholds for gene and transcript counts. Data were log-normalized NormalizeData(), and highly variable genes were identified using variance-stabilizing transformation (VST) FindVariableFeatures(). The data were then scaled ScaleData() and subjected to Principal Component Analysis (PCA) RunPCA(). The top 30 principal components (PCs) were used for Louvain clustering (FindClusters(), resolution = 0.5) and nearest-neighbor graph construction FindNeighbors(). For visualization, Uniform Manifold Approximation and Projection (UMAP) was applied RunUMAP(), and clusters were visualized with DimPlot(). The final Seurat objects were saved as .rds files for downstream analyses.

### 6. Correlation, MI, and matrix distance methods

#### 3.1 hdWGCNA

hdWGCNA was conducted using the “hdWGCNA” R package to identify gene co-expression modules in single-cell RNA-seq data across four Arabidopsis samples: WER, CORTEX, PET111, and SCR. Seurat objects were first prepared by including the top highly variable genes along with promoter-reporter and native gene pairs. Metacells were constructed using the MetacellsByGroups() function, which aggregated transcriptomic profiles based on cell type annotations to reduce noise and improve co-expression signal robustness. Each metacell expression matrix was normalized via NormalizeMetacells(). The expression matrix was set using SetDatExpr(), specifying relevant cell type groups for module detection. Network topology was assessed by testing a range of soft thresholding powers using TestSoftPowers(), and optimal values were selected based on scale-free topology criteria. A signed gene co-expression network was then constructed using ConstructNetwork(), and topological overlap matrices (TOMs) were used to rank gene-gene co-expression relationships. The module membership of genes was retrieved using GetModules(), and dendrograms were generated using PlotDendrogram() for each sample. The co-expression strength between each reporter gene and its native counterpart was evaluated based on TOM values and module co-membership.

#### 3.2 CSCORE

Gene-gene correlation analysis was performed using the CS-CORE method, implemented via the “CSCORE” R package, installed from GitHub. Normalized single-cell expression matrices were generated from Seurat objects using the NormalizeData() function. Highly variable genes were identified using the variance-stabilizing transformation (VST) method through FindVariableFeatures(), with the top 1,000/2,000 variable genes selected for analysis. The expression matrix was then scaled using ScaleData(). CS-CORE was applied to the scaled data using the CSCORE() function, with the selected variable features provided as input. The resulting gene-gene correlation estimates were extracted and ranked to evaluate co-expression between promoter-reporter genes and their corresponding native genes.

#### 3.3 Pearson, Spearman, and Kendall

Pairwise correlations between genes were computed using Pearson, Spearman, and Kendall correlation coefficients. For each gene pair, correlation tests were performed using the cor.test() function in R across three data layers extracted from Seurat objects: raw counts, log-normalized data, and scaled data, corresponding to the "counts", "data", and "scale.data" assay layers. Correlation analyses were conducted across three subsets of cells: (1) all cells, (2) cells expressing at least one of the two genes, and (3) cells expressing both genes. A cell was considered to express a gene if it had a non-zero count for that gene in the raw count matrix.

#### 3.4 Biweight midcorrelation (bicor)

Bicor was computed using the bicor() function from the “WGCNA” R package to assess gene-gene relationships. Similar to Pearson, Spearman, and Kendall methods, bicor was applied to three data layers extracted from Seurat objects and evaluated across three cell subsets.

#### 3.5 Mutual information

Mutual information between gene pairs was estimated by first discretizing gene expression values into five bins using equal-frequency binning via the discretize() function from the “infotheo” R package. Similar to the Pearson, Spearman, and Kendall methods, mutual information analysis was applied to three data layers extracted from Seurat objects and conducted across three cell subsets.

#### 3.6 p-sctransform

Gene expression data were normalized using the SCTransform method from the “sctransform” R package. SCTransform was applied to Seurat objects using SCTransform(), with residual features set to include highly variable genes along with the gene pair of interest. Pairwise gene correlations were then computed using the Pearson method on the scaled expression values in the SCT assay slot.

#### 3.7 Euclidean, Manhattan, and Chebyshev distances

Euclidean, Manhattan, and Chebyshev distances were computed based on absolute differences in gene expression across cells, with Euclidean defined as the square root of the sum of squared differences, Manhattan as the sum of absolute differences, and Chebyshev as the maximum absolute difference.

### 4. Simulation with scDesign2

Simulated data were generated using the scDesign2 package with a fitted Gaussian copula model to capture gene-gene dependencies and negative binomial marginals to model count distributions. The original count matrix was extracted from a Seurat object. Pairwise gene-gene correlations were computed using Pearson and Spearman methods for both the original and simulated count matrices. To focus on highly expressed genes, the top 200 genes were selected based on their mean expression across all cells. Correlation matrices were computed on this subset, and hierarchical clustering was performed on the Pearson matrix from the original data to determine a consistent gene ordering for visualization.

### 5. Pseudo-bulk analysis

Cells were grouped by cluster identities, and the mean expression and 95% confidence intervals, extracted from the raw count matrix, were calculated for each gene across clusters. Pairwise Pearson, Spearman, and Kendall correlations were then computed between the cluster-level mean expression values of the two genes.

### 6. Imputation methods

#### 6.1 scImpute

The “scImpute” R package was used to perform gene expression imputation on a subset of 2,000 genes, including highly variable genes identified from the Seurat object and genes of interest extracted from the raw count matrix. The scimpute() function was applied with a dropout probability threshold of 0.5, an estimated number of cell clusters (Kcluster = 10), and 10 cores for parallel processing. The imputed expression matrix was then used to compute pairwise Pearson and Spearman correlations between the genes of interest to evaluate co-expression recovery.

#### 6.2 SAVER

The “SAVER” R package was used to perform gene expression imputation on a subset of genes, including the top 500 highly variable genes from the Seurat object and the genes of interest. The saver() function was applied with pred.genes.only = TRUE to impute only the selected genes. The posterior mean estimates of gene expression were extracted from the $estimate slot and rounded using the round() function for downstream analysis.

#### 6.3 AE

An Autoencoder neural network was implemented in PyTorch to impute gene expression values in single-cell RNA-seq data. Input data were log-transformed (log1p) and trained using a feedforward Autoencoder consisting of an encoder (input -> 400 -> 200 units) and decoder (200 -> 400 -> input units) with LeakyReLU activations. The model was trained using mean squared error (MSE) loss and optimized with the Adam optimizer. The training was conducted over a user-defined number of epochs, with the loss history recorded and visualized. After training, the model was used to reconstruct the input data, which were then inverse transformed (expm1) and rounded before saving. Pearson and Spearman correlations were later computed on the imputed data to evaluate co-expression recovery.

#### 6.4 VAE

A Variational Autoencoder (VAE) was implemented in PyTorch to impute gene expression values in single-cell RNA-seq data. Input data were log-transformed (log1p) and used to train a VAE consisting of an encoder, reparameterization module, and decoder. The model was trained using a combined loss function including mean squared error (MSE) for reconstruction and Kullback-Leibler divergence (KLD) for latent regularization. After training, reconstructed expression values were inverse transformed (expm1) and rounded before saving.

#### 6.5 GAN

A Generative Adversarial Network (GAN) was implemented in PyTorch to perform gene expression imputation on single-cell RNA-seq data. Input data were log-transformed (log1p) and used to train a GAN consisting of a Generator for data reconstruction and a Discriminator for distinguishing real from synthetic samples. The Generator was trained using mean squared error (MSE) loss to minimize reconstruction error, while the Discriminator used binary cross-entropy loss (BCE) to classify real and generated data. The model was trained over multiple epochs with separate learning rates for the Generator and Discriminator. After training, imputed expression values were inverse transformed (expm1) and rounded before saving.

### 7. TF-target genes validate biological

Pairwise correlations among target genes of transcription factors (TFs) were computed in both original and imputed single-cell RNA-seq datasets. For each dataset, the expression profiles of each TF’s target genes were extracted, and the mean expression level across all target genes was calculated. To assess regulatory coherence, pairwise correlations among the target genes of each TF were calculated using Pearson, Spearman, or Kendall methods. The resulting correlation values were then aggregated by computing the average correlation coefficient per TF, enabling a comparative evaluation of gene-gene relationships between the original and imputed data. TF-target gene relationships were obtained from ConnecTF^34^.

### 8. qPCR

Cells were isolated from Arabidopsis root tissues and sorted into GFP-positive and GFP-negative populations using fluorescence-activated cell sorting (FACS), based on the expression of marker lines PET111, WER, SCR, and CORTEX. Total RNA was extracted from the sorted cells using the RNeasy Plus Micro Kit (Qiagen), following the manufacturer’s protocol. Genomic DNA contamination was eliminated by on-column DNase I digestion. Complementary DNA (cDNA) synthesis was performed using the iScript cDNA Synthesis Kit (Bio-Rad). Quantitative PCR (qPCR) was carried out using the iTaq Universal SYBR Green Supermix (Bio-Rad) on an ABI 7500 Fast Real-Time PCR System (Applied Biosystems). B_Actin was used as the reference gene for normalization. Each reaction was run in technical triplicates, and primer sequences are available upon request.

## Code Availability

The scripts for the analysis can be found at out GitHub repository (https://github.com/ct-tranchau/scCoBench)

## Data Availability

The data are deposited in GEO database (GSE295703)

## Supplementary information

Supplementary Figures 1-11

## Acknowledgements

Step 2.1 of Figure 1 was created with BioRender (https://www.biorender.com).

## Peer review information

## Authors’ contributions

S.L. supervised the whole project. T.N.C. analyzed the data. K.H.R. & J.S. generated single-cell RNA-seq data. R.A. & B.O.R.B conducted the wet-bench validation. All the authors wrote the manuscript. All authors read and approved the final manuscript.

## Declarations

### Ethics approval and consent to participate

Not applicable.

### Consent for publication

Not applicable.

