## Supplementary_Figures for "scCoBench: Benchmarking single cell RNA-seq co-expression using promoter-reporter lines"

### Slide 1
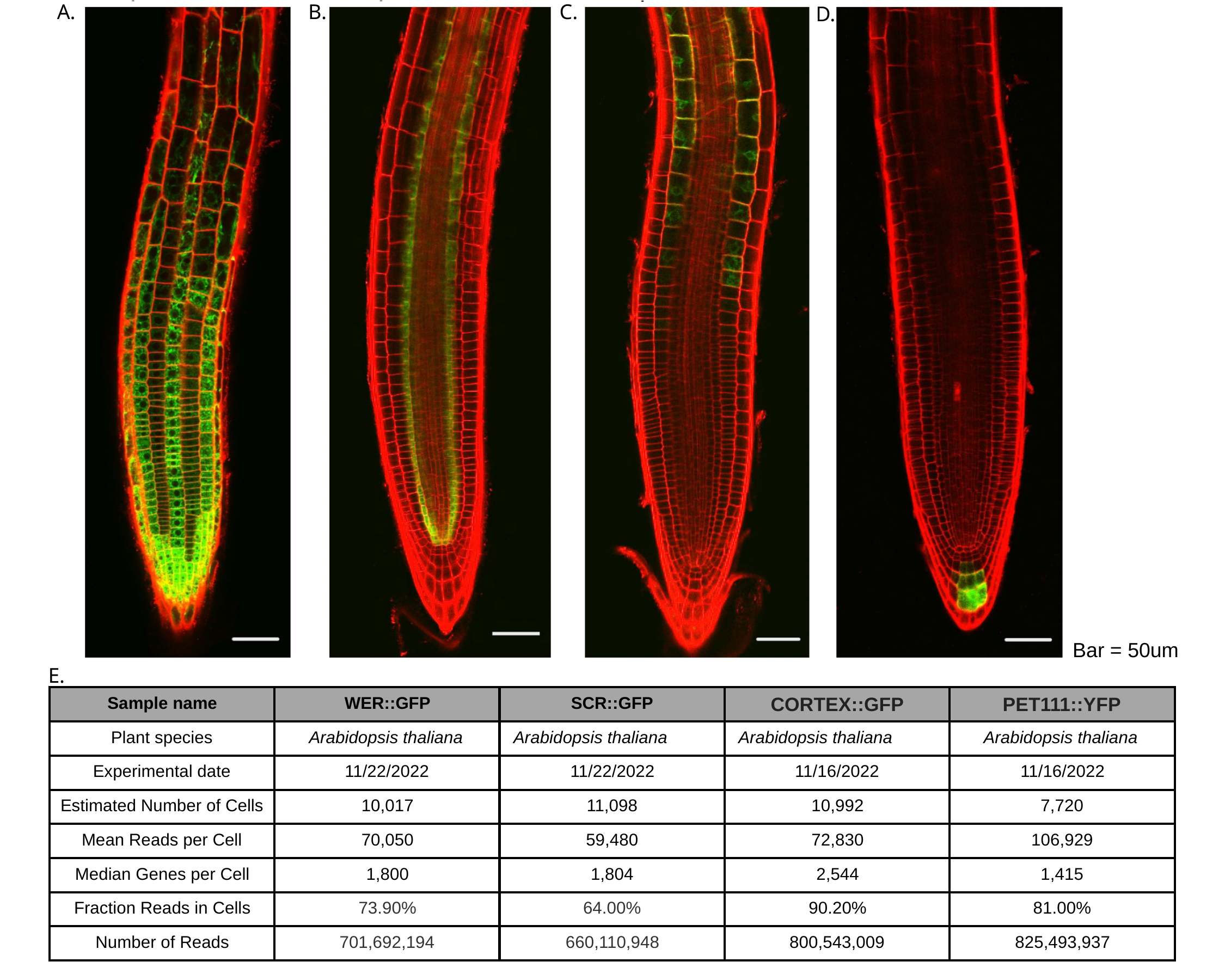

A.
B.
C.
D.
Bar = 50um
E.
| Sample name | WER::GFP | SCR::GFP | CORTEX::GFP | PET111::YFP |
| --- | --- | --- | --- | --- |
| Plant species | Arabidopsis thaliana | Arabidopsis thaliana | Arabidopsis thaliana | Arabidopsis thaliana |
| Experimental date | 11/22/2022 | 11/22/2022 | 11/16/2022 | 11/16/2022 |
| Estimated Number of Cells | 10,017 | 11,098 | 10,992 | 7,720 |
| Mean Reads per Cell | 70,050 | 59,480 | 72,830 | 106,929 |
| Median Genes per Cell | 1,800 | 1,804 | 2,544 | 1,415 |
| Fraction Reads in Cells | 73.90% | 64.00% | 90.20% | 81.00% |
| Number of Reads | 701,692,194 | 660,110,948 | 800,543,009 | 825,493,937 |

### Slide 2
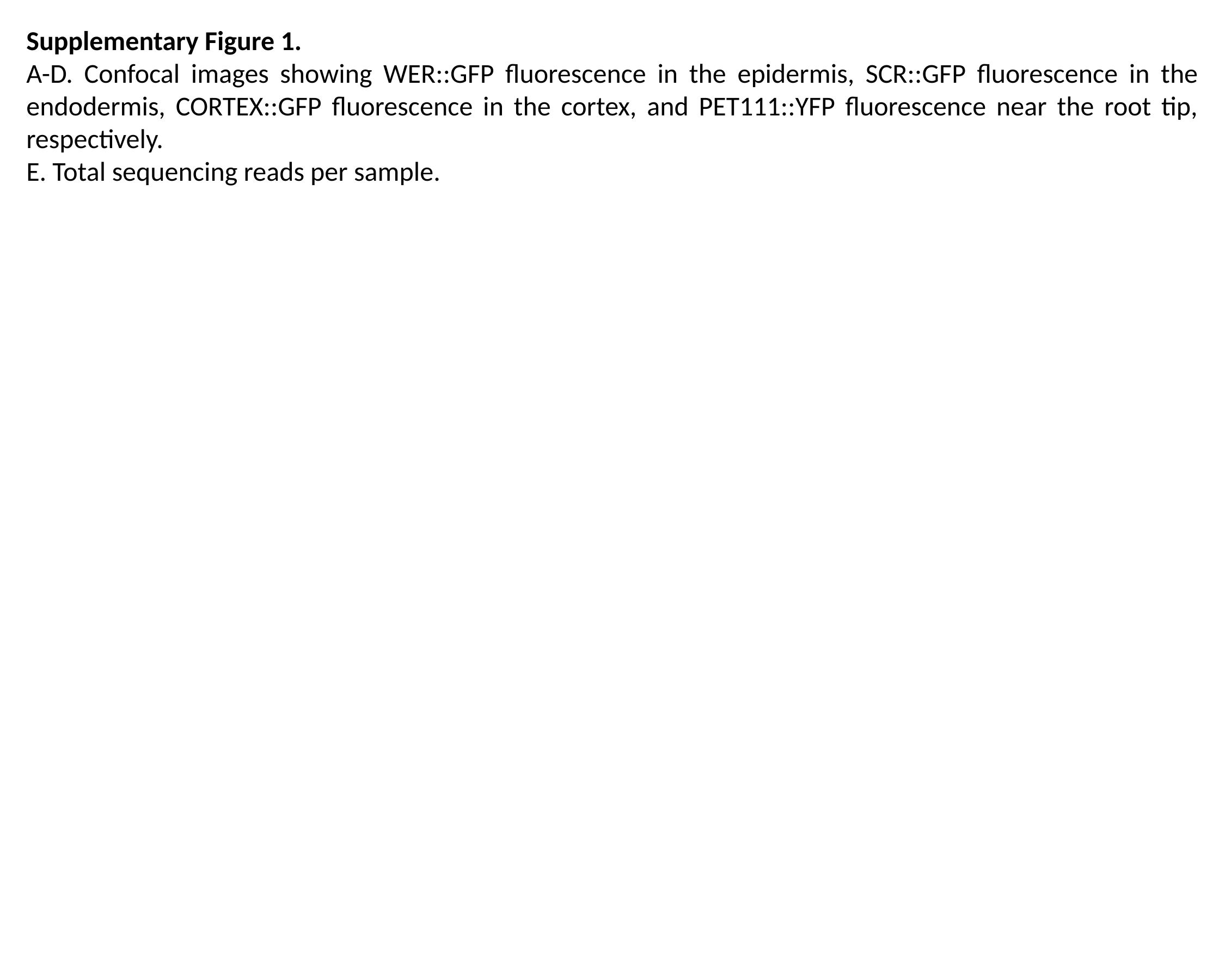

Supplementary Figure 1.
A-D. Confocal images showing WER::GFP fluorescence in the epidermis, SCR::GFP fluorescence in the endodermis, CORTEX::GFP fluorescence in the cortex, and PET111::YFP fluorescence near the root tip, respectively.
E. Total sequencing reads per sample.

### Slide 3
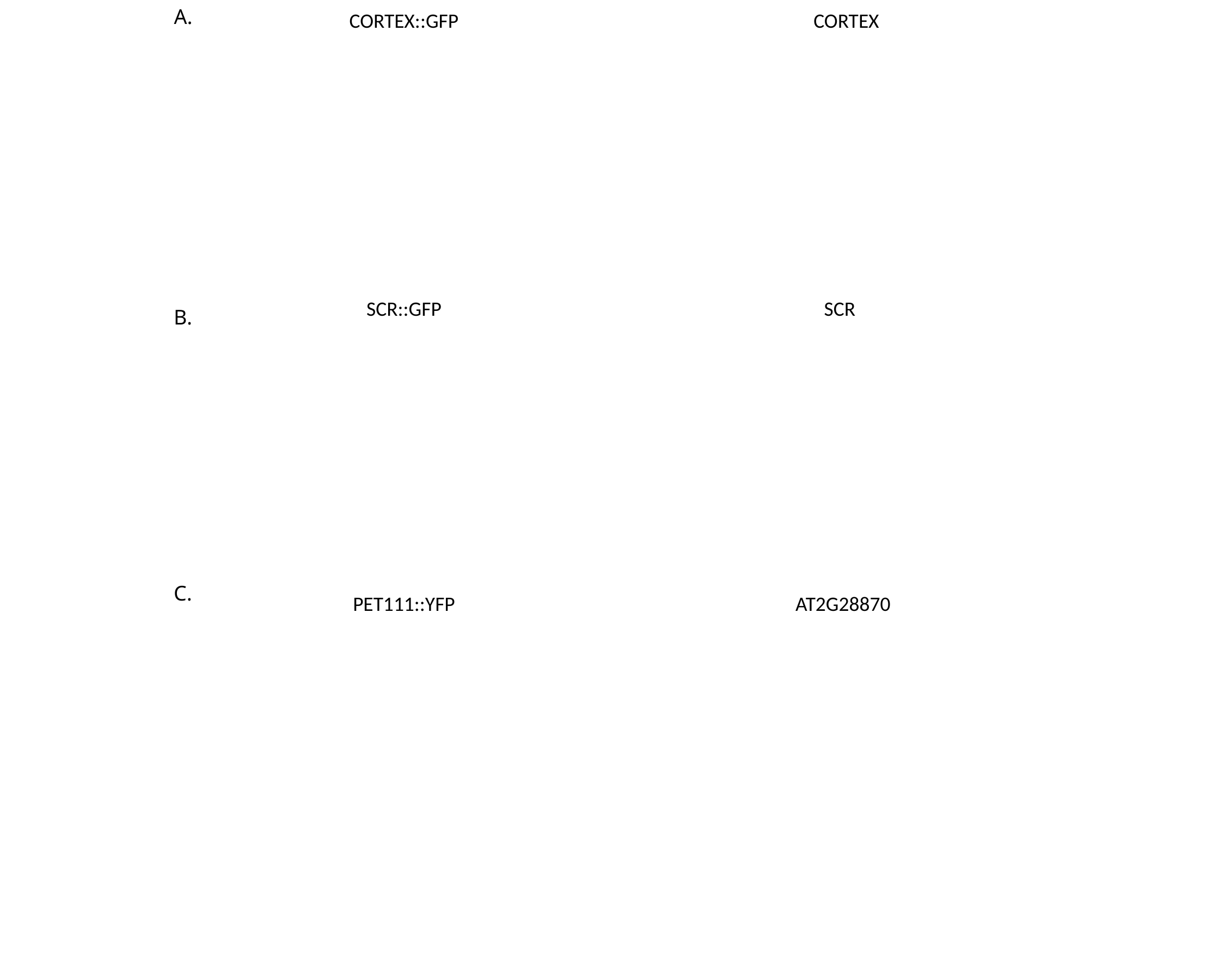

A.
CORTEX::GFP
CORTEX
SCR::GFP
SCR
B.
C.
PET111::YFP
AT2G28870

### Slide 4
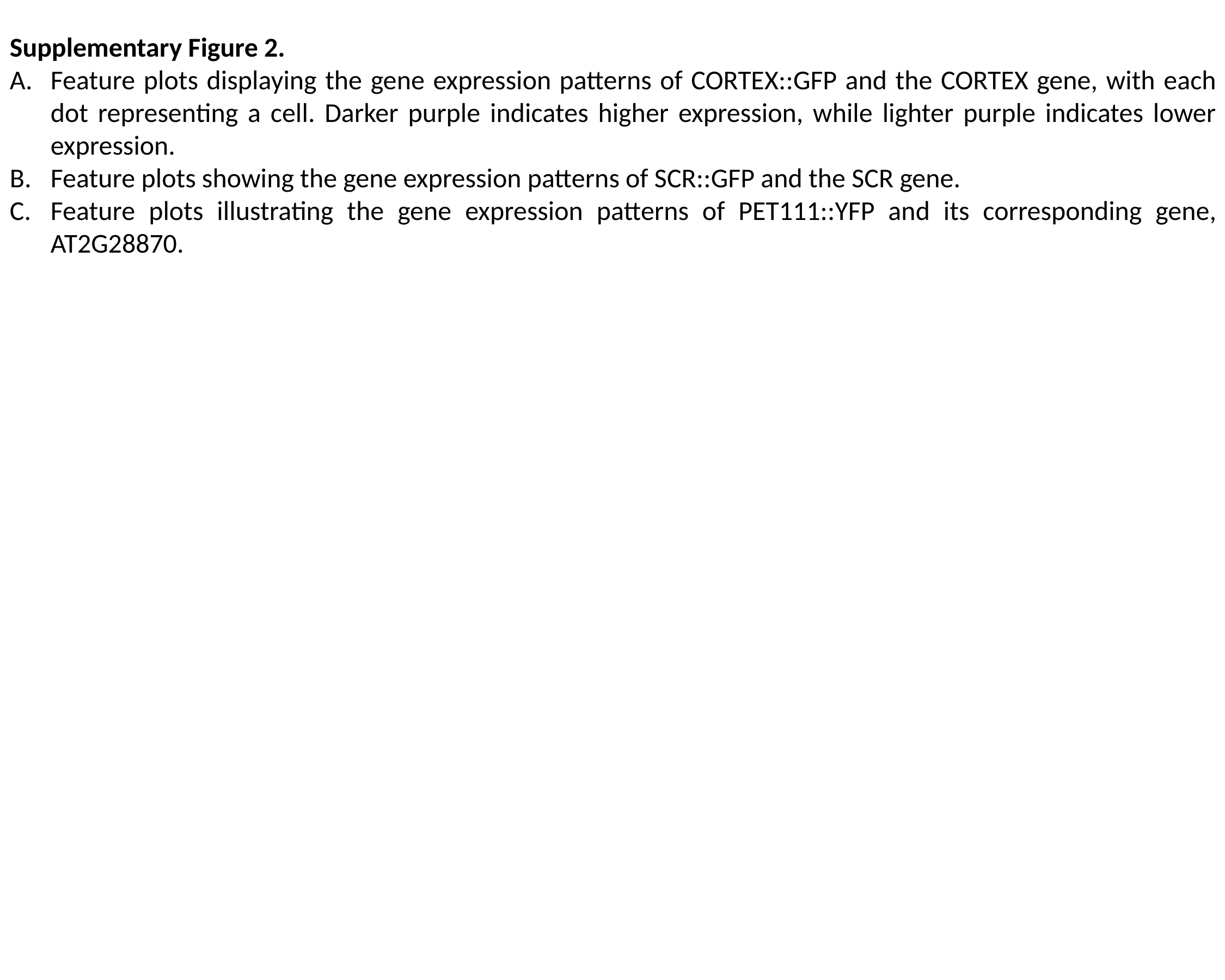

Supplementary Figure 2.
Feature plots displaying the gene expression patterns of CORTEX::GFP and the CORTEX gene, with each dot representing a cell. Darker purple indicates higher expression, while lighter purple indicates lower expression.
Feature plots showing the gene expression patterns of SCR::GFP and the SCR gene.
Feature plots illustrating the gene expression patterns of PET111::YFP and its corresponding gene, AT2G28870.

### Slide 5
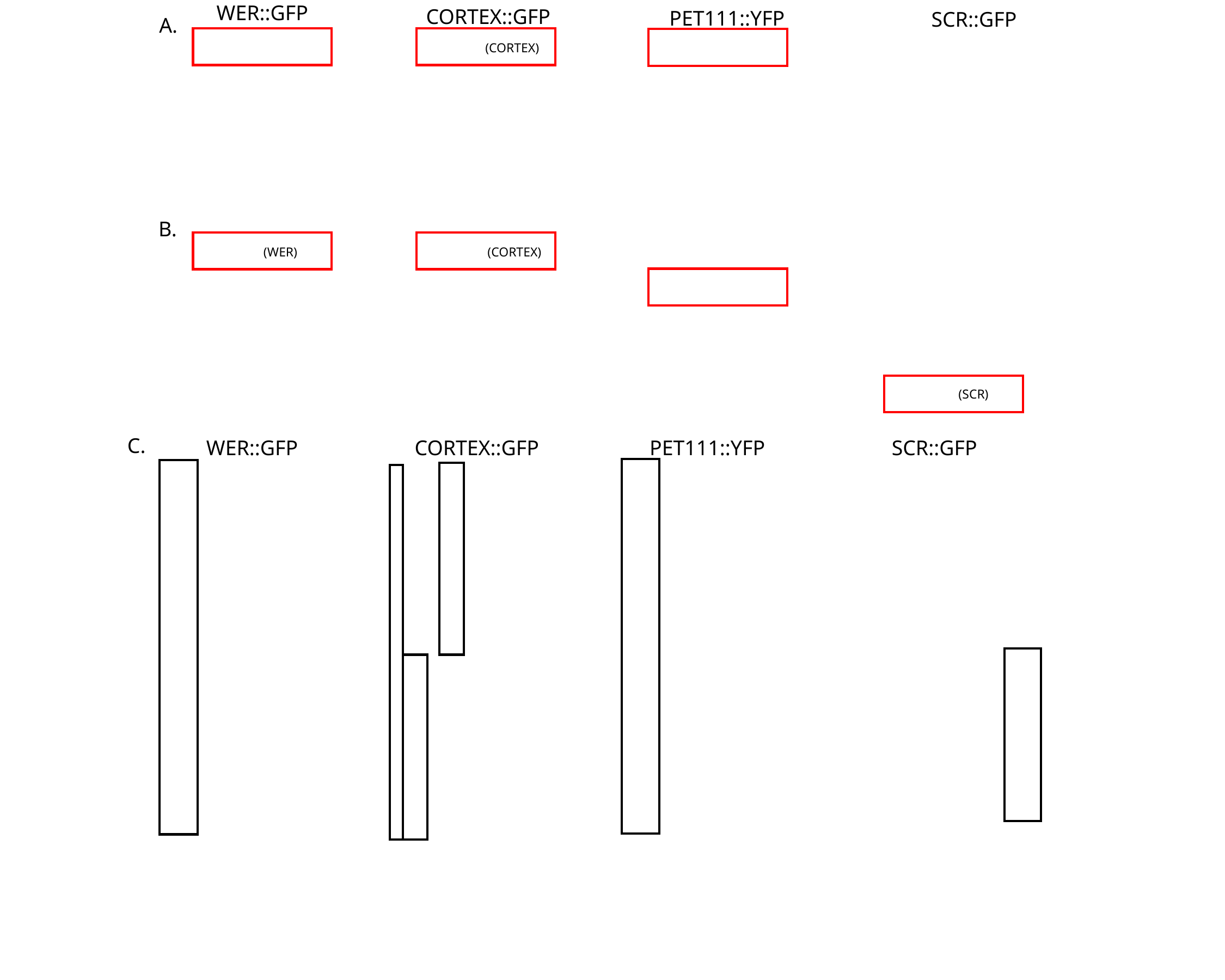

WER::GFP
CORTEX::GFP
PET111::YFP
SCR::GFP
A.
(WER)
(CORTEX)
B.
(WER)
(CORTEX)
(SCR)
C.
WER::GFP
CORTEX::GFP
PET111::YFP
SCR::GFP

### Slide 6
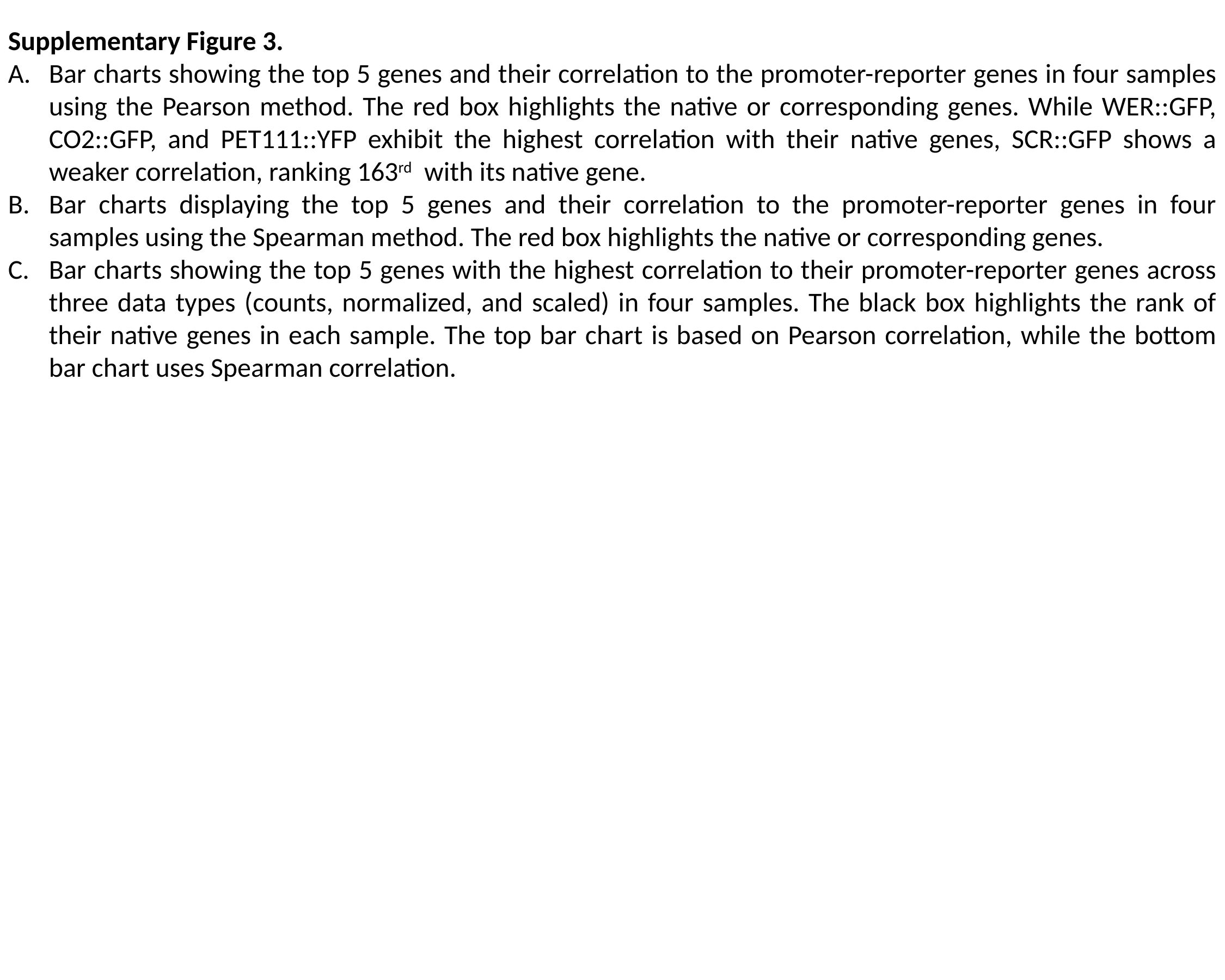

Supplementary Figure 3.
Bar charts showing the top 5 genes and their correlation to the promoter-reporter genes in four samples using the Pearson method. The red box highlights the native or corresponding genes. While WER::GFP, CO2::GFP, and PET111::YFP exhibit the highest correlation with their native genes, SCR::GFP shows a weaker correlation, ranking 163^rd^ with its native gene.
Bar charts displaying the top 5 genes and their correlation to the promoter-reporter genes in four samples using the Spearman method. The red box highlights the native or corresponding genes.
Bar charts showing the top 5 genes with the highest correlation to their promoter-reporter genes across three data types (counts, normalized, and scaled) in four samples. The black box highlights the rank of their native genes in each sample. The top bar chart is based on Pearson correlation, while the bottom bar chart uses Spearman correlation.

### Slide 7
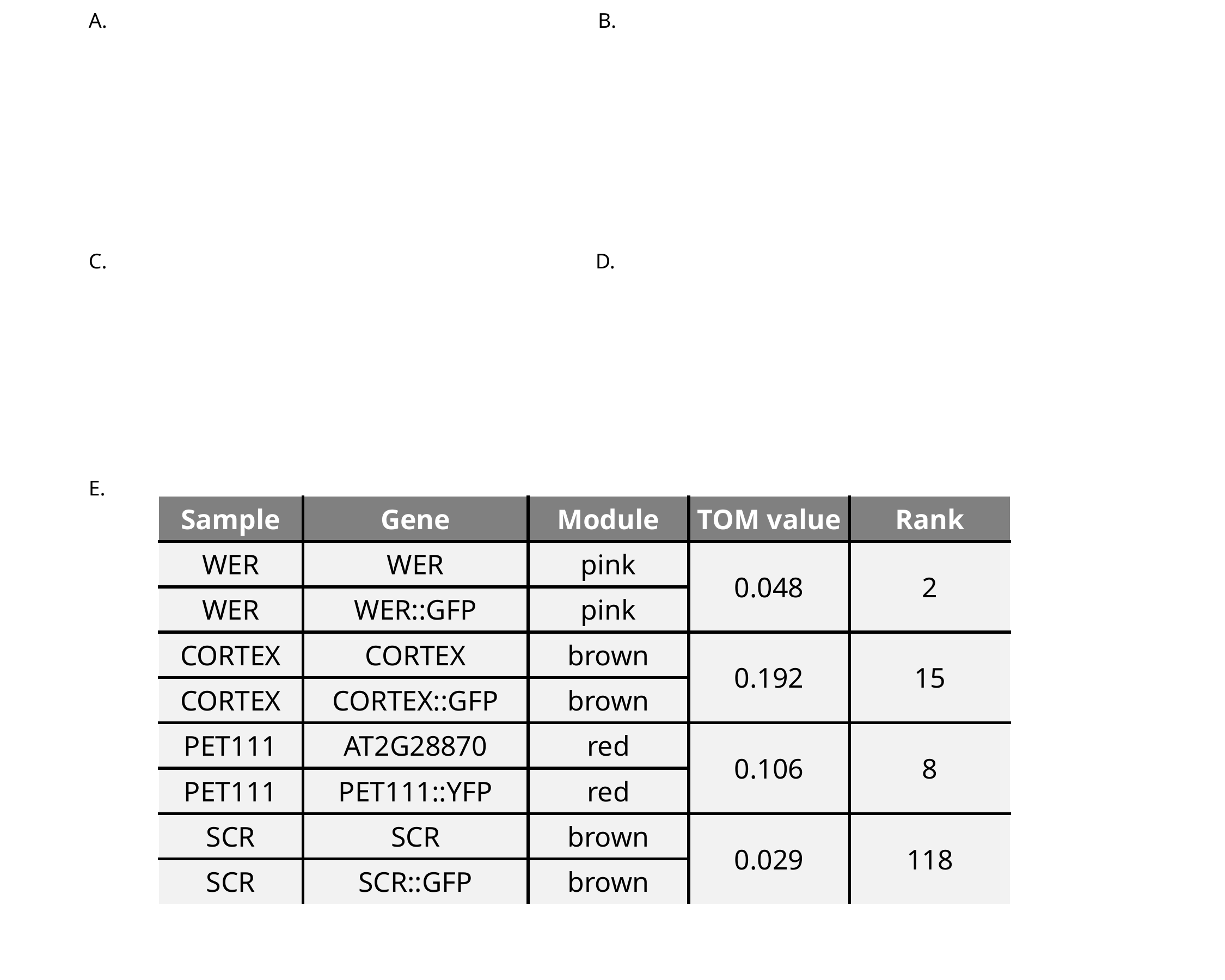

A.
B.
C.
D.
E.
| Sample | Gene | Module | TOM value | Rank |
| --- | --- | --- | --- | --- |
| WER | WER | pink | 0.048 | 2 |
| WER | WER::GFP | pink | | |
| CORTEX | CORTEX | brown | 0.192 | 15 |
| CORTEX | CORTEX::GFP | brown | | |
| PET111 | AT2G28870 | red | 0.106 | 8 |
| PET111 | PET111::YFP | red | | |
| SCR | SCR | brown | 0.029 | 118 |
| SCR | SCR::GFP | brown | | |

### Slide 8
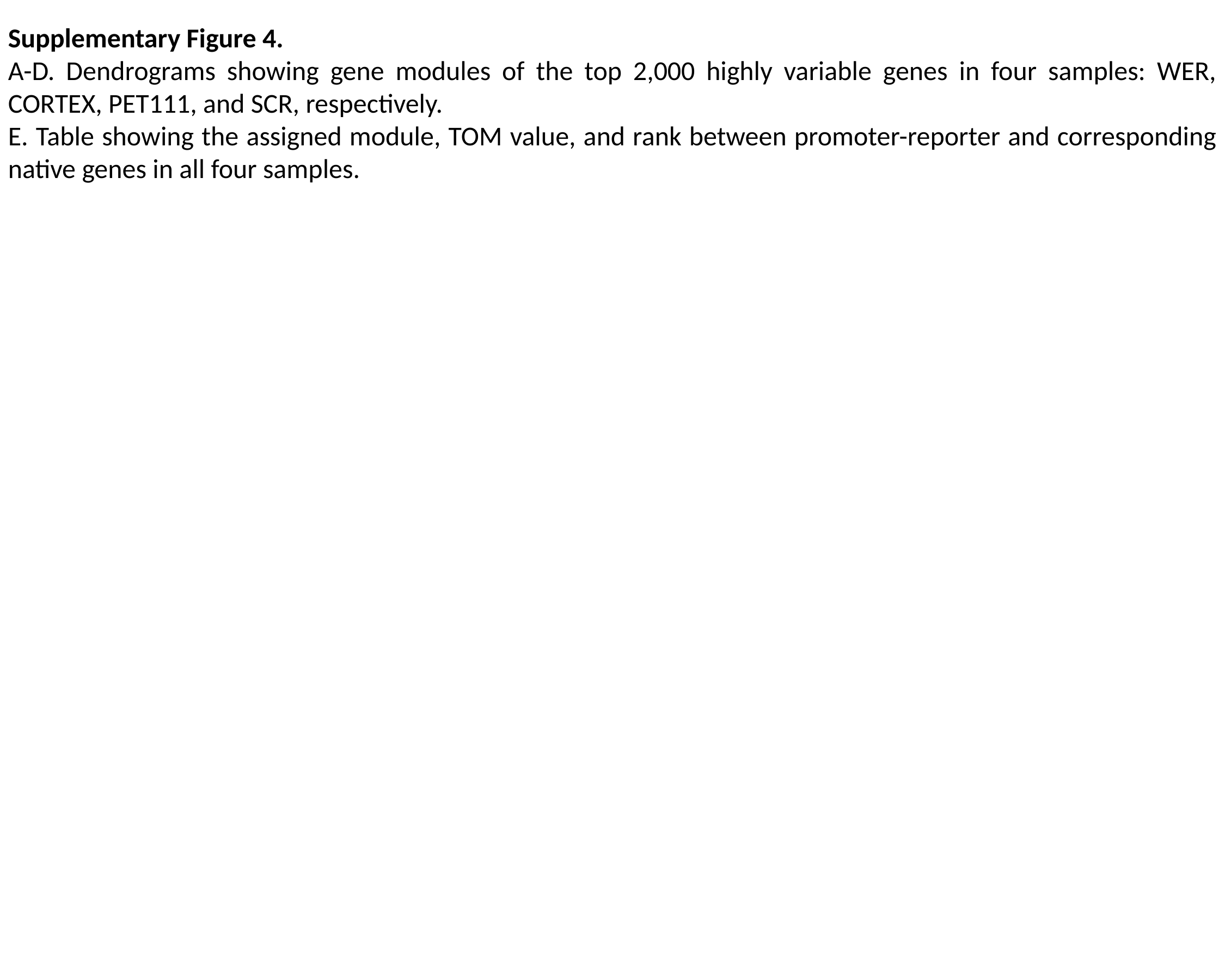

Supplementary Figure 4.
A-D. Dendrograms showing gene modules of the top 2,000 highly variable genes in four samples: WER, CORTEX, PET111, and SCR, respectively.
E. Table showing the assigned module, TOM value, and rank between promoter-reporter and corresponding native genes in all four samples.

### Slide 9
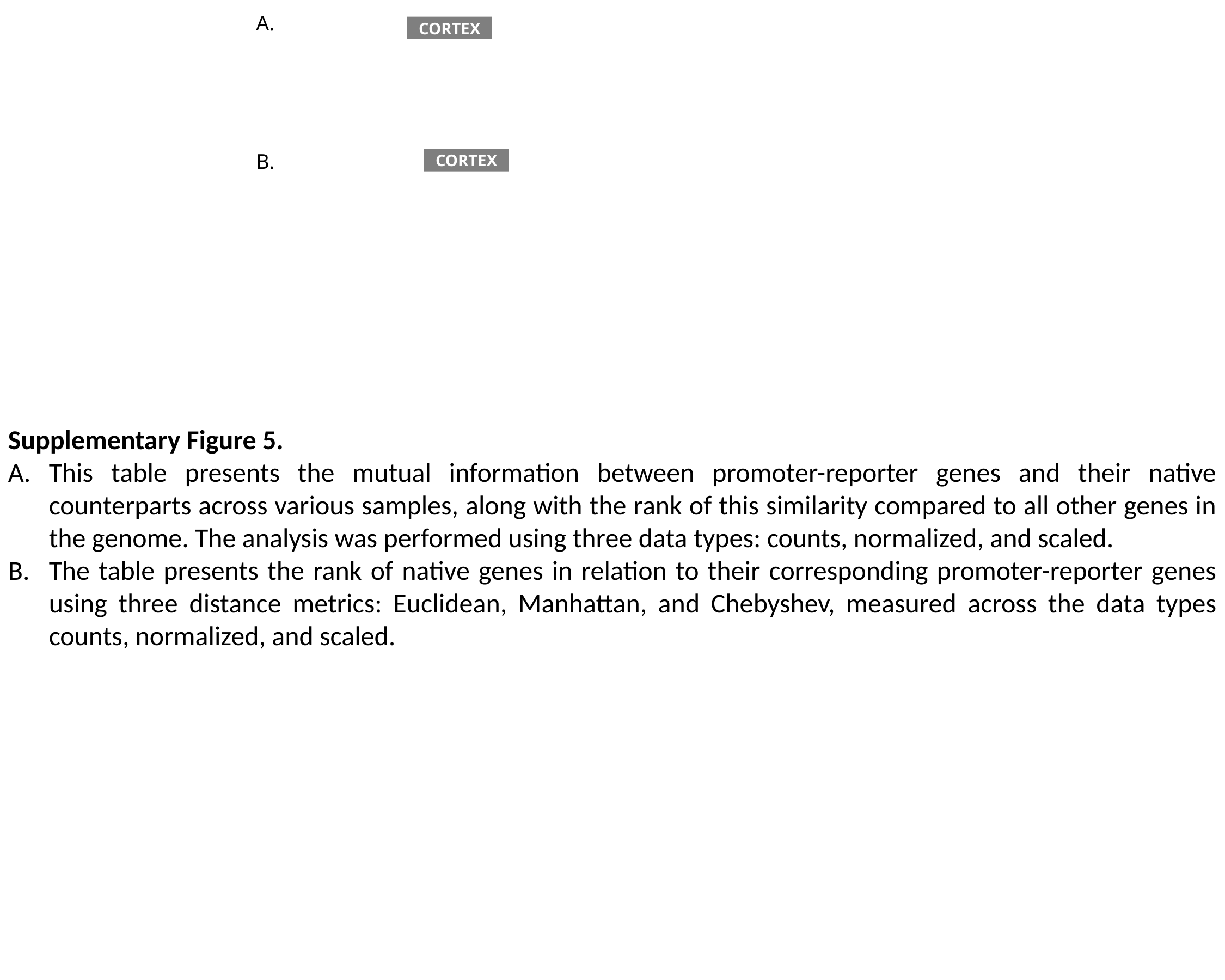

A.
CORTEX
B.
CORTEX
Supplementary Figure 5.
This table presents the mutual information between promoter-reporter genes and their native counterparts across various samples, along with the rank of this similarity compared to all other genes in the genome. The analysis was performed using three data types: counts, normalized, and scaled.
The table presents the rank of native genes in relation to their corresponding promoter-reporter genes using three distance metrics: Euclidean, Manhattan, and Chebyshev, measured across the data types counts, normalized, and scaled.

### Slide 10
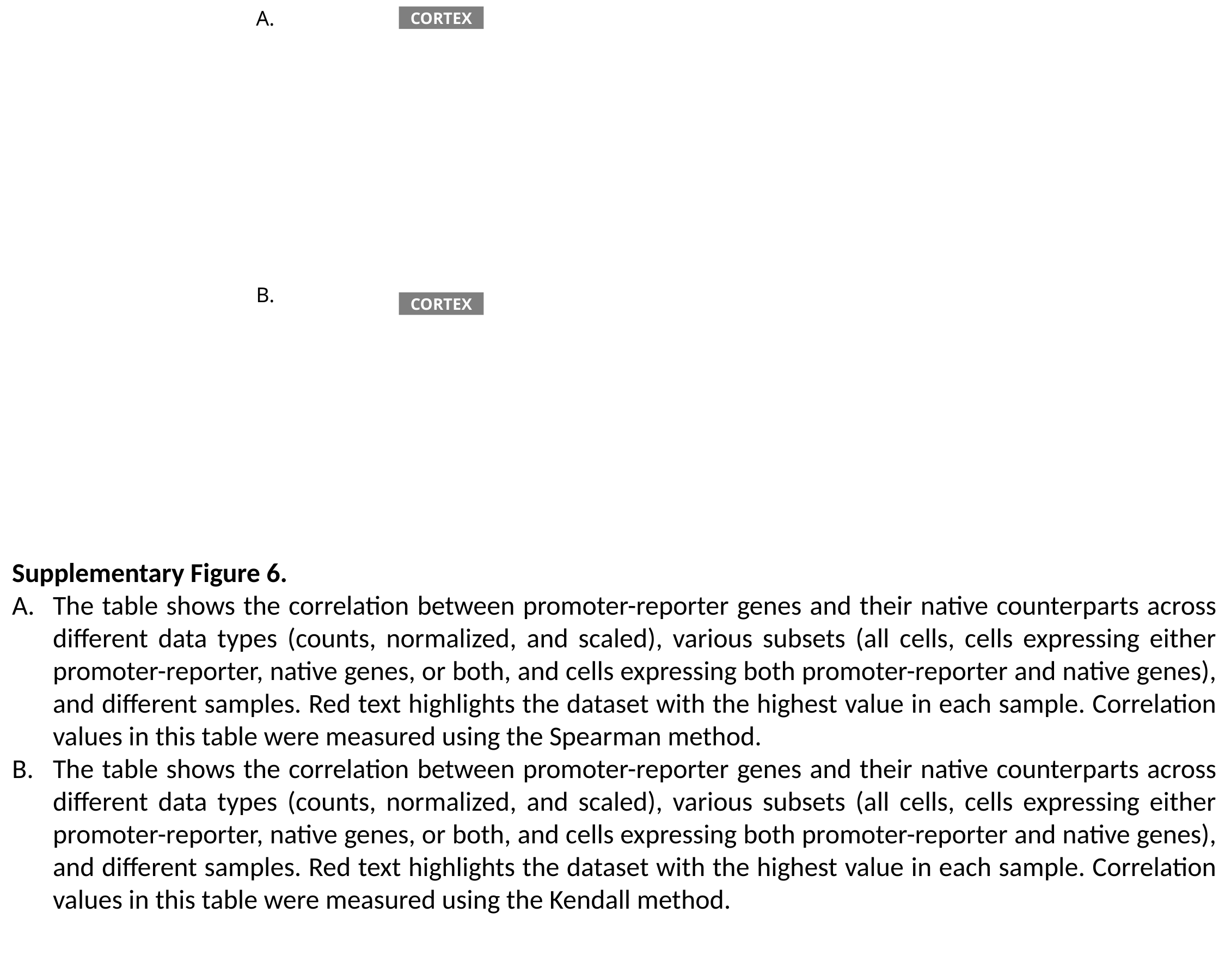

A.
CORTEX
B.
CORTEX
Supplementary Figure 6.
The table shows the correlation between promoter-reporter genes and their native counterparts across different data types (counts, normalized, and scaled), various subsets (all cells, cells expressing either promoter-reporter, native genes, or both, and cells expressing both promoter-reporter and native genes), and different samples. Red text highlights the dataset with the highest value in each sample. Correlation values in this table were measured using the Spearman method.
The table shows the correlation between promoter-reporter genes and their native counterparts across different data types (counts, normalized, and scaled), various subsets (all cells, cells expressing either promoter-reporter, native genes, or both, and cells expressing both promoter-reporter and native genes), and different samples. Red text highlights the dataset with the highest value in each sample. Correlation values in this table were measured using the Kendall method.

### Slide 11
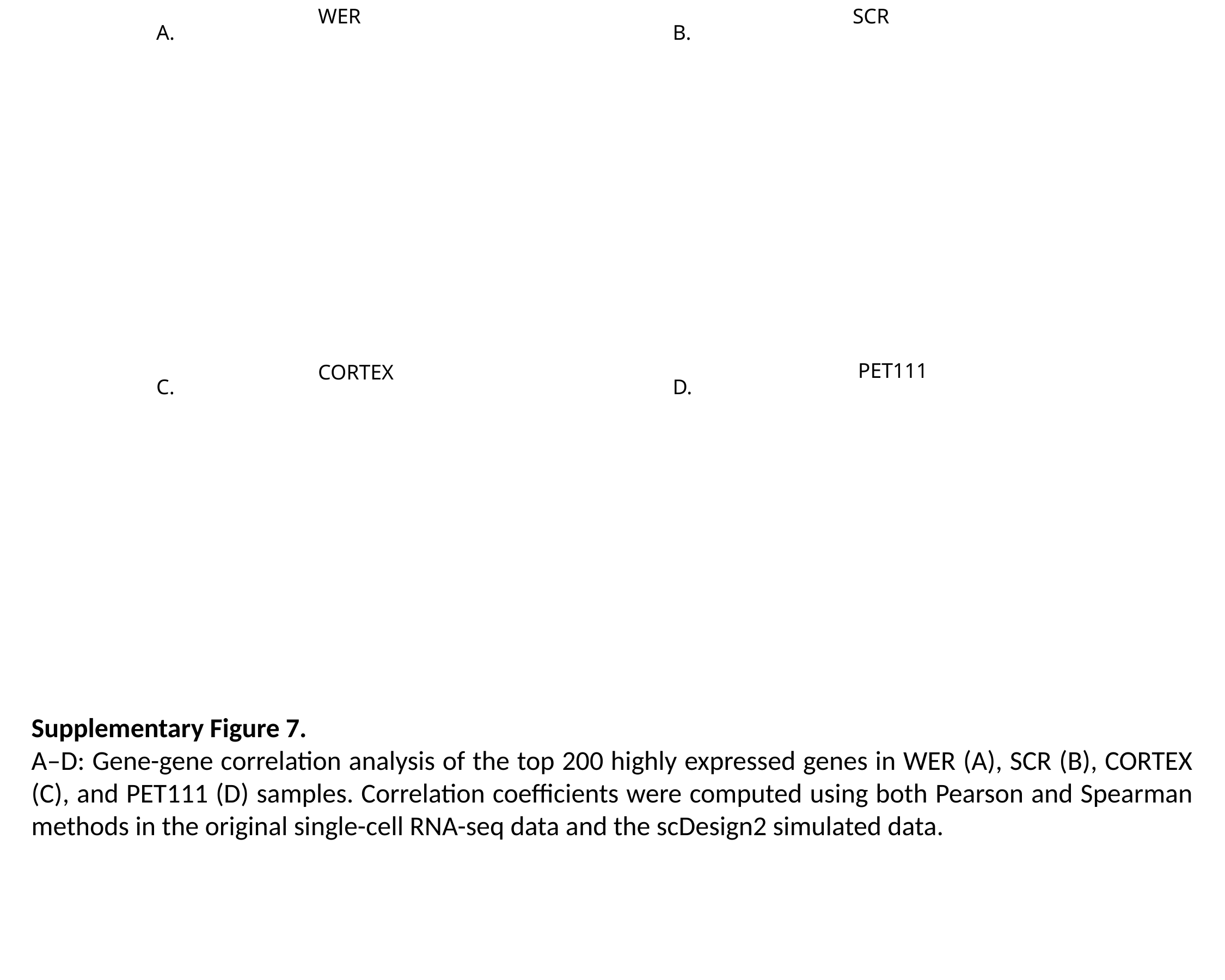

WER
SCR
A.
B.
PET111
CORTEX
C.
D.
Supplementary Figure 7.
A–D: Gene-gene correlation analysis of the top 200 highly expressed genes in WER (A), SCR (B), CORTEX (C), and PET111 (D) samples. Correlation coefficients were computed using both Pearson and Spearman methods in the original single-cell RNA-seq data and the scDesign2 simulated data.

### Slide 12
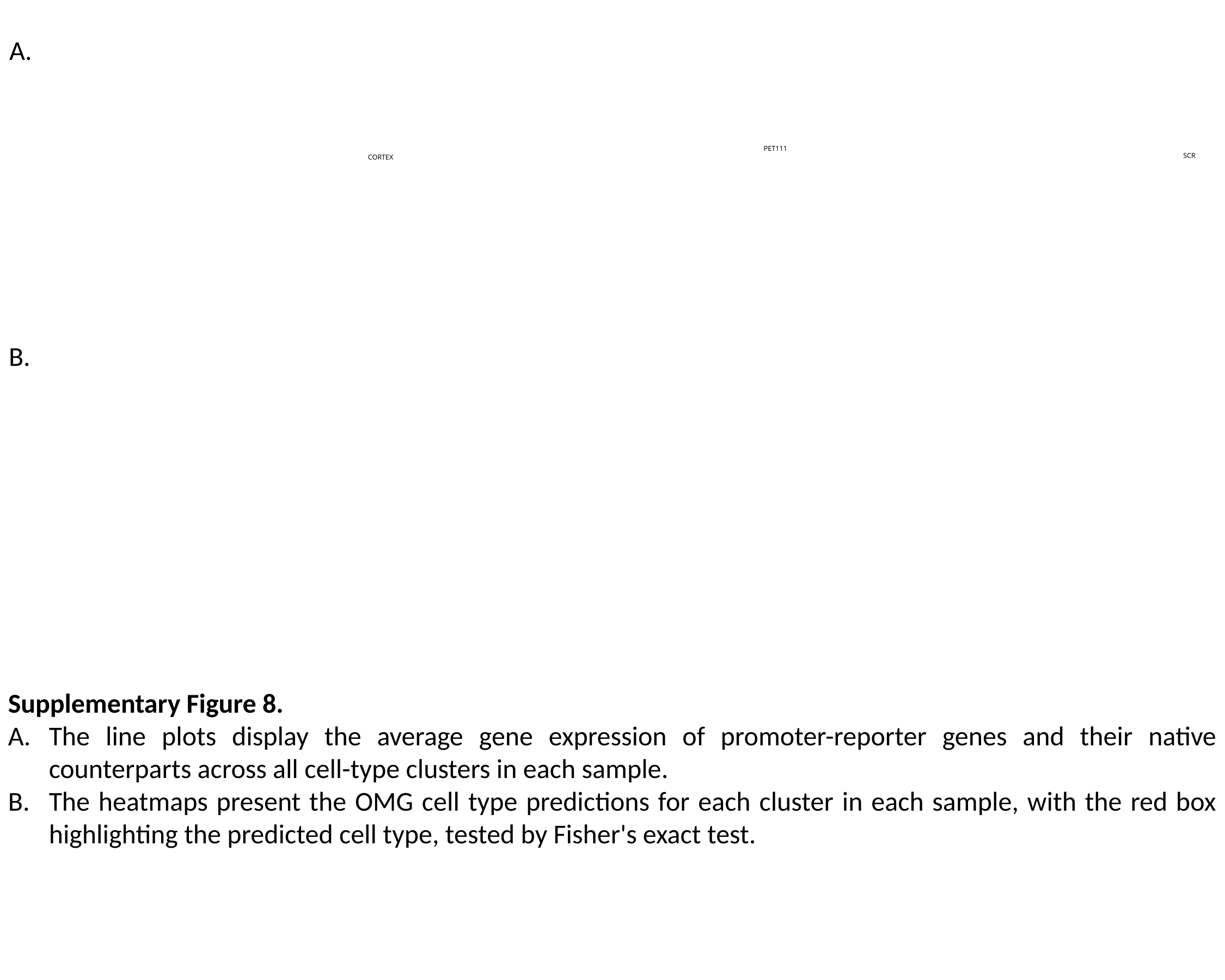

A.
PET111
SCR
CORTEX
B.
Supplementary Figure 8.
The line plots display the average gene expression of promoter-reporter genes and their native counterparts across all cell-type clusters in each sample.
The heatmaps present the OMG cell type predictions for each cluster in each sample, with the red box highlighting the predicted cell type, tested by Fisher's exact test.

### Slide 13
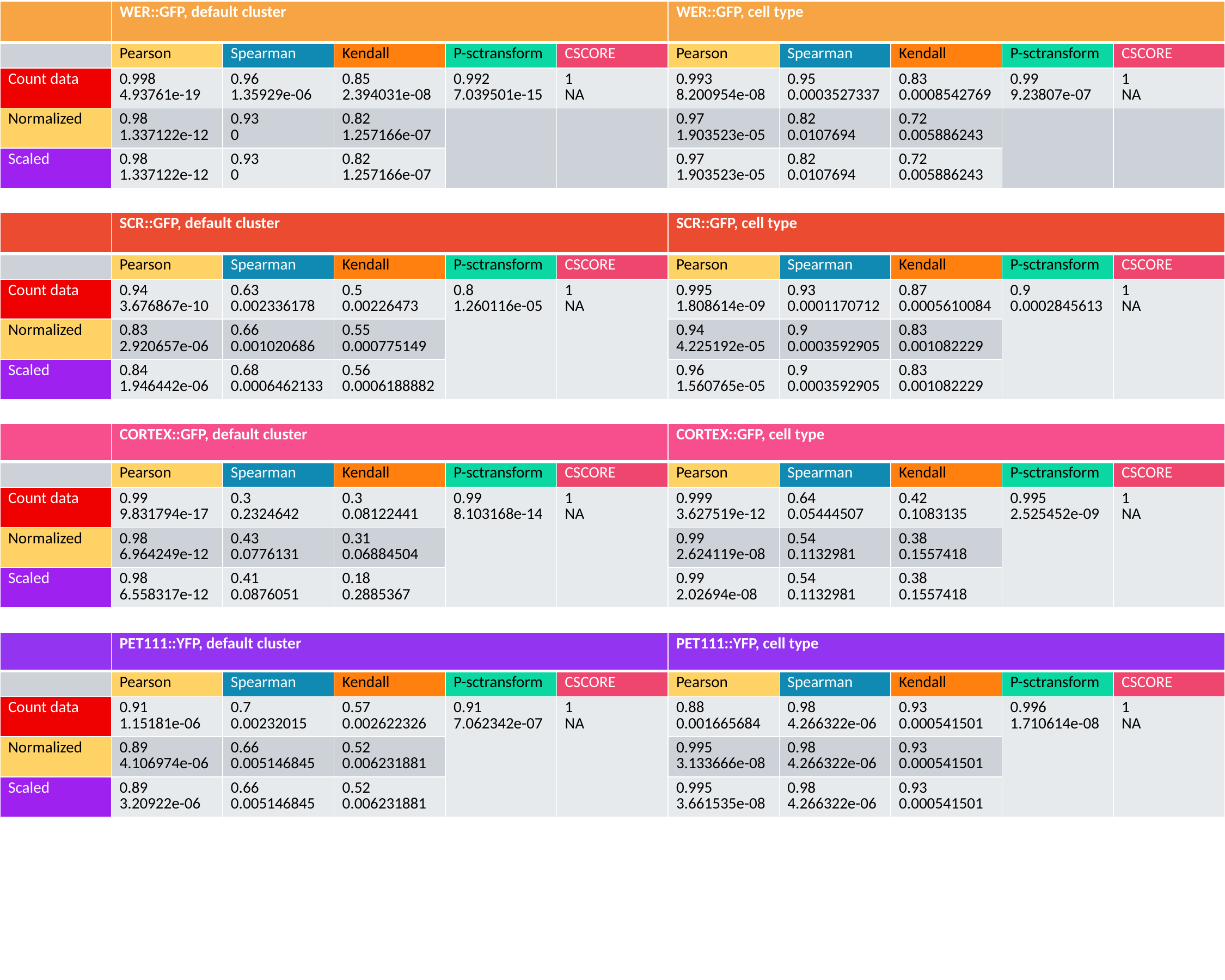

| | WER::GFP, default cluster | | | | | WER::GFP, cell type | | | | |
| --- | --- | --- | --- | --- | --- | --- | --- | --- | --- | --- |
| | Pearson | Spearman | Kendall | P-sctransform | CSCORE | Pearson | Spearman | Kendall | P-sctransform | CSCORE |
| Count data | 0.998 4.93761e-19 | 0.96 1.35929e-06 | 0.85 2.394031e-08 | 0.992 7.039501e-15 | 1 NA | 0.993 8.200954e-08 | 0.95 0.0003527337 | 0.83 0.0008542769 | 0.99 9.23807e-07 | 1 NA |
| Normalized | 0.98 1.337122e-12 | 0.93 0 | 0.82 1.257166e-07 | | | 0.97 1.903523e-05 | 0.82 0.0107694 | 0.72 0.005886243 | | |
| Scaled | 0.98 1.337122e-12 | 0.93 0 | 0.82 1.257166e-07 | | | 0.97 1.903523e-05 | 0.82 0.0107694 | 0.72 0.005886243 | | |
| | SCR::GFP, default cluster | | | | | SCR::GFP, cell type | | | | |
| --- | --- | --- | --- | --- | --- | --- | --- | --- | --- | --- |
| | Pearson | Spearman | Kendall | P-sctransform | CSCORE | Pearson | Spearman | Kendall | P-sctransform | CSCORE |
| Count data | 0.94 3.676867e-10 | 0.63 0.002336178 | 0.5 0.00226473 | 0.8 1.260116e-05 | 1 NA | 0.995 1.808614e-09 | 0.93 0.0001170712 | 0.87 0.0005610084 | 0.9 0.0002845613 | 1 NA |
| Normalized | 0.83 2.920657e-06 | 0.66 0.001020686 | 0.55 0.000775149 | | | 0.94 4.225192e-05 | 0.9 0.0003592905 | 0.83 0.001082229 | | |
| Scaled | 0.84 1.946442e-06 | 0.68 0.0006462133 | 0.56 0.0006188882 | | | 0.96 1.560765e-05 | 0.9 0.0003592905 | 0.83 0.001082229 | | |
| | CORTEX::GFP, default cluster | | | | | CORTEX::GFP, cell type | | | | |
| --- | --- | --- | --- | --- | --- | --- | --- | --- | --- | --- |
| | Pearson | Spearman | Kendall | P-sctransform | CSCORE | Pearson | Spearman | Kendall | P-sctransform | CSCORE |
| Count data | 0.99 9.831794e-17 | 0.3 0.2324642 | 0.3 0.08122441 | 0.99 8.103168e-14 | 1 NA | 0.999 3.627519e-12 | 0.64 0.05444507 | 0.42 0.1083135 | 0.995 2.525452e-09 | 1 NA |
| Normalized | 0.98 6.964249e-12 | 0.43 0.0776131 | 0.31 0.06884504 | | | 0.99 2.624119e-08 | 0.54 0.1132981 | 0.38 0.1557418 | | |
| Scaled | 0.98 6.558317e-12 | 0.41 0.0876051 | 0.18 0.2885367 | | | 0.99 2.02694e-08 | 0.54 0.1132981 | 0.38 0.1557418 | | |
| | PET111::YFP, default cluster | | | | | PET111::YFP, cell type | | | | |
| --- | --- | --- | --- | --- | --- | --- | --- | --- | --- | --- |
| | Pearson | Spearman | Kendall | P-sctransform | CSCORE | Pearson | Spearman | Kendall | P-sctransform | CSCORE |
| Count data | 0.91 1.15181e-06 | 0.7 0.00232015 | 0.57 0.002622326 | 0.91 7.062342e-07 | 1 NA | 0.88 0.001665684 | 0.98 4.266322e-06 | 0.93 0.000541501 | 0.996 1.710614e-08 | 1 NA |
| Normalized | 0.89 4.106974e-06 | 0.66 0.005146845 | 0.52 0.006231881 | | | 0.995 3.133666e-08 | 0.98 4.266322e-06 | 0.93 0.000541501 | | |
| Scaled | 0.89 3.20922e-06 | 0.66 0.005146845 | 0.52 0.006231881 | | | 0.995 3.661535e-08 | 0.98 4.266322e-06 | 0.93 0.000541501 | | |

### Slide 14
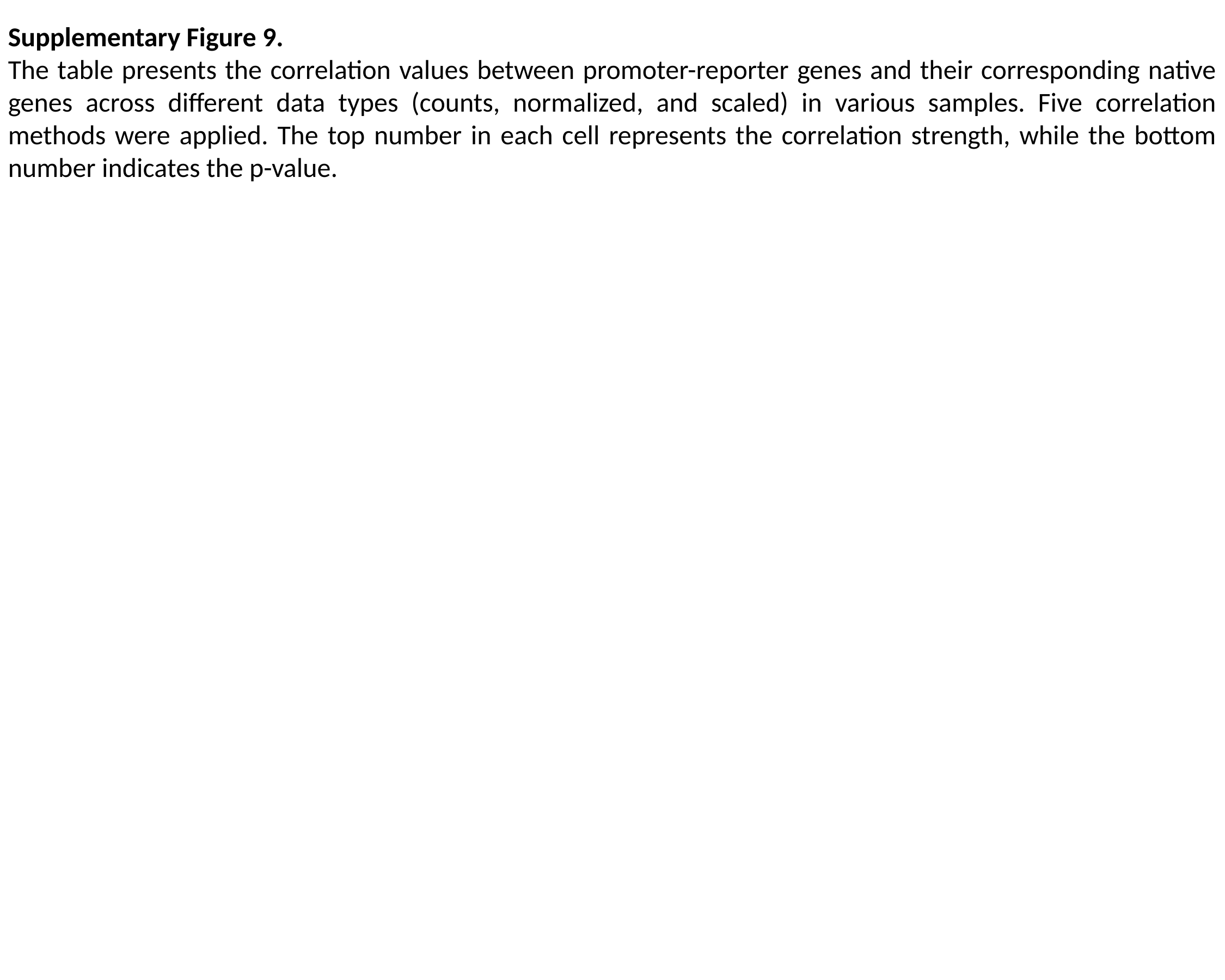

Supplementary Figure 9.
The table presents the correlation values between promoter-reporter genes and their corresponding native genes across different data types (counts, normalized, and scaled) in various samples. Five correlation methods were applied. The top number in each cell represents the correlation strength, while the bottom number indicates the p-value.

### Slide 15
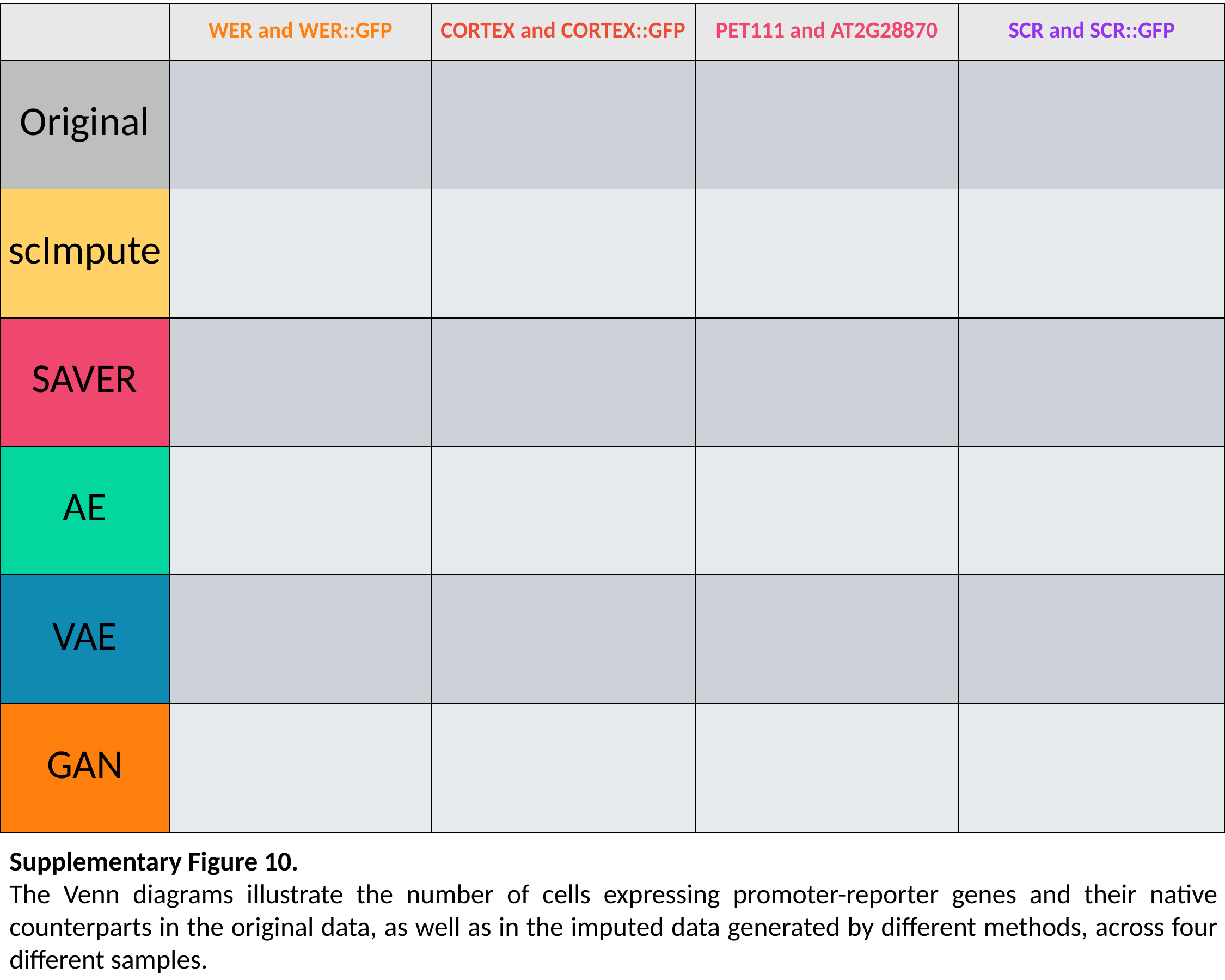

| | WER and WER::GFP | CORTEX and CORTEX::GFP | PET111 and AT2G28870 | SCR and SCR::GFP |
| --- | --- | --- | --- | --- |
| Original | | | | |
| scImpute | | | | |
| SAVER | | | | |
| AE | | | | |
| VAE | | | | |
| GAN | | | | |
Supplementary Figure 10.
The Venn diagrams illustrate the number of cells expressing promoter-reporter genes and their native counterparts in the original data, as well as in the imputed data generated by different methods, across four different samples.

### Slide 16
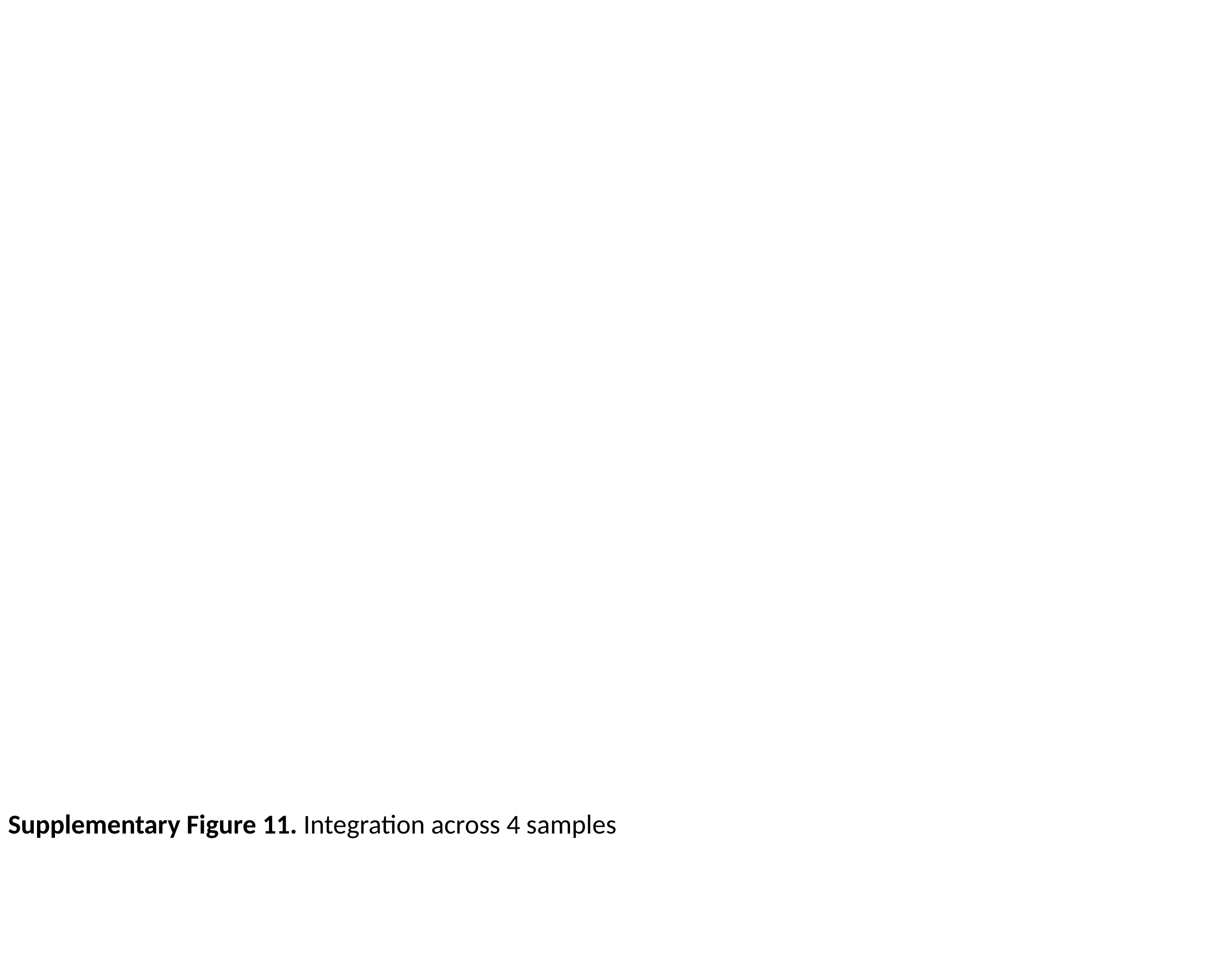

Supplementary Figure 11. Integration across 4 samples
